# EMT network-based feature selection improves prognosis prediction in lung adenocarcinoma

**DOI:** 10.1101/410472

**Authors:** Borong Shao, Maria M Bjaanæs, Helland Åslaug, Christof Schütte, Tim Conrad

## Abstract

Various feature selection algorithms have been proposed to identify cancer prognostic biomarkers. In recent years, however, their reproducibility is criticized. The performance of feature selection algorithms is shown to be affected by the datasets, underlying networks and evaluation metrics. One of the causes is the curse of dimensionality, which makes it hard to select the features that generalize well on independent data. Even the integration of biological networks does not mitigate this issue because the networks are large and many of their components are not relevant for the phenotype of interest. With the availability of multi-omics data, integrative approaches are being developed to build more robust predictive models. In this scenario, the higher data dimensions create greater challenges.

We proposed a phenotype relevant network-based feature selection (PRNFS) framework and demonstrated its advantages in lung cancer prognosis prediction. We constructed cancer prognosis relevant networks based on epithelial mesenchymal transition (EMT) and integrated them with different types of omics data for feature selection. With less than 2.5% of the total dimensionality, we obtained EMT prognostic signatures that achieved remarkable prediction performance (average AUC values *>*0.8), very significant sample stratifications, and meaningful biological interpretations. In addition to finding EMT signatures from different omics data levels, we combined these single-omics signatures into multi-omics signatures, which improved sample stratifications significantly. Both single- and multi-omics EMT signatures were tested on independent multi-omics lung cancer datasets and significant sample stratifications were obtained.

## Introduction

Prognosis prediction is necessary for cancer clinical decision making. Traditionally, cancer prognosis prediction is based on clinical variables such as tumor stage, age, and disease history, where the information of a patient is compared against population caner registries [1]. However, these clinical parameters are insufficient to accurately predict the risk of patients [2] as histologically similar tumors can be of completely different diseases at the molecular level [3, 4]. Therefore, molecular signatures are needed to give more accurate prognosis predictions. Nowadays we can obtain tumor molecular profiles in greater details. As one of the large scale projects, the Cancer Genome Atlas (TCGA) provides access to genomic, transcriptomic, epigenomic, and proteomic data from more than 11,000 cases in 33 cancer types and subtypes [5]. Using these data, researchers aim to build better prognosis prediction models. This has been found as very challenging due to the high dimensionality of omics data, where the number of features far exceeds the number of samples. This is often addressed as *the curse of dimensionality*. When the data lie in high dimensions, the samples become very sparse. This can cause the lack of statistical significance and over-fitting of machine learning models. Fortunately, not all features are relevant for predicting the phenotype of interest. It is desired to find the molecular signatures that capture the footprint of the phenotype so that the signatures can be employed on unseen samples.

Various feature selection methods were proposed to find molecular signatures. Early reviews categorized them into three categories: filter, wrapper, and embedded methods [6, 7]. Although many important algorithms were introduced, network-based feature selection algorithms were not included. After reviewing the literature, we found three main categories of network-based feature selection algorithms. The first category involves network-guided search. An algorithm identifies subnetworks that can best differentiate different phenotype groups. Each subnetwork is aggregated to produce one feature (called metagene) and eventually the metagenes are used as features for training predictive models. Different scoring functions were proposed to rank the subnetworks. For example, [8] used mutual information as the scoring function and the addition operator to aggregate subnetworks. [9, 10] used the p-value of Cox PH model in defining the scoring functions. [11] dichotomized features and defined a scoring function based on information theory. [12] tested the effects of different aggregation operators on the prediction performance.

The second category of methods uses network-based regularization. Regularization methods such as Lasso have been widely applied for feature selection. To integrate network information, the penalty term takes into account the network connectivity. Adjacency matrix *A* and Laplacian matrix *L* are frequently used to represent a network *G* to be included in the penalty term. The majority of methods in this category are based on linear classifiers and can be written in the following form:

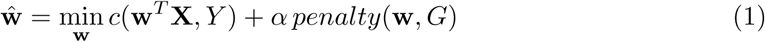

LFor example, in graph Lasso *penalty*(**w**, *G*) = *λ*║w║_1_ + (1 – *λ*)Ʃ_*i,j*_ *A*_*i*_,_*j*_(**w**_*i*_ − **w**_*j*_). This forces adjacent nodes to have similar weights [13, 14]. Using similar formulations, [15] proposed a network-constrained regularization and feature selection method on genomic data. [16] added a network regularization term to the log-likelihood function of the Cox proportional hazard model. [17] developed a network-constrained support vector machine algorithm, where the network-based regularization term is added to the objective function of SVM.

The third category of methods involves iterative updates of node importance scores. Frequently used algorithms include network propagation and random walk. [18, 19] adapted Google’s PageRank algorithm to rank genes in a network. Genes are assigned initial ranks **r**^[**0**]^ *∈* ℝ^*N*^. Then the rank of each gene is updated iteratively depending on the ranks of genes that are linked to it. For gene *j*, its rank from 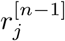 or 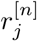 is updated as:

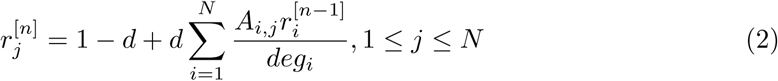

where *deg*_*i*_ is the degree of the *i*th gene and *d* is a fixed parameter. By iterating until convergence a gene will be highly ranked if it is linked to other highly ranked genes. [20] used random walk kernel to smooth gene-wise t-statistics over the network. This is achieved by assigning each node an initial score based on t-test and then multiplying it with the random walk kernel. The *p*-step random walk kernel is used as a similarity measure to capture the relatedness of two nodes in the network. It is defined as:

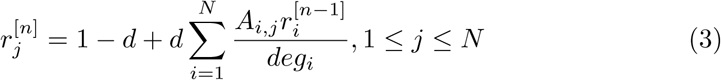

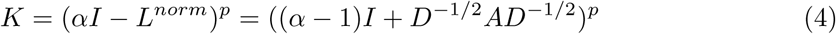

where *L*^*norm*^ is the normalized graph Laplacian matrix, *α* is a constant, and *p* is the number of random walk steps. The network-smoothed t-statistic 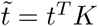 is used to measure node importance. Similarly, random walk-based scoring of network components is applied in [21] to prioritize functional networks.

Besides algorithmic difference, various biological networks have been employed in network-based feature selection algorithms. A list of these molecular interactions is given in Table 1. In these studies, it was shown that superior features could be selected by the integration of networks. However, recent studies showed that network-based feature selection methods did not significantly improve prediction performance, but mainly contributed to the biological interpretations of the signatures [22–24]. [22] compared 14 feature selection methods, 8 of which integrated network information and 6 of which did not, on 6 breast cancer datasets with respect to their prediction accuracy, signature stability and biological interpretations. The results showed that network-based features in most cases could not improve prediction accuracy significantly. [23, 24] showed that when a correction of feature set size was performed, the stability of network-based features was not higher than single features.

**Table 1.**
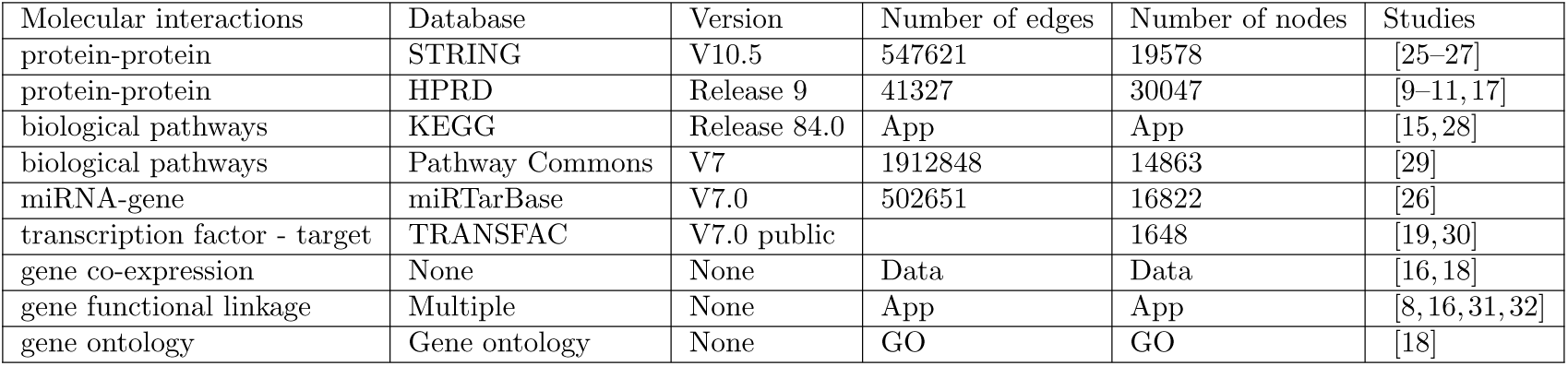
Frequently used molecular/gene interaction networks in network-based feature selection studies. We listed below the basic information of the networks as well as exemplary studies that employed the networks. With STRING database we only considered the edges with confidence scores *≥* 0.9. When a database has information of many species, only Homo sapiens was considered. In the 4th and 5th columns, *Data* means that the network size is dependent on the dimensions of data. *App* means that the network size is dependent on the application. *GO* means that the network size is dependent on gene ontology terms.

| Molecular interactions | Database | Version | Number of edges | Number of nodes | Studies |
| --- | --- | --- | --- | --- | --- |
| protein-protein | STRING | V10.5 | 547621 | 19578 | [25–27] |
| protein-protein | HPRD | Release 9 | 41327 | 30047 | [9–11, 17] |
| biological pathways | KEGG | Release 84.0 | App | App | [15, 28] |
| biological pathways | Pathway Commons | V7 | 1912848 | 14863 | [29] |
| miRNA-gene | miRTarBase | V7.0 | 502651 | 16822 | [26] |
| transcription factor - target | TRANSFAC | V7.0 public |  | 1648 | [19, 30] |
| gene co-expression | None | None | Data | Data | [16, 18] |
| gene functional linkage | Multiple | None | App | App | [8, 16, 31, 32] |
| gene ontology | Gene ontology | None | GO | GO | [18] |

Regardless of whether network information is integrated, finding robust molecular signatures is a challenging task. Due to the high data dimensions, it is easy to find a feature subset that fits the training data very well but hard to have good generalization. Studies showed that there was hardly considerable overlap among biomarkers identified in different studies for the same disease [33–35]. Even taking random feature sets gave comparable prediction performance [36]. The existence of many feature subsets that perform similarly well on the training set makes it difficult to identify the true signatures. Note that the randomness of signatures is also observed when network information is integrated. [23] tested different network-based feature selection algorithms on six breast cancer datasets in prognosis prediction. They showed that the randomization of network structure, which destroyed biological information, did not deteriorate the prediction performance of the selected features. [24] extended the experiments in [23] by comparing more prognosis signatures. In the end, similar results were observed.

We suppose that the main reason for these counter-intuitive results is the curse of dimensionality, where selecting molecular signatures is hard given the limited amount of samples. In principle, molecular signatures should give better predictions than random features, because it is shown in biological research that certain genes are supposed to be more important than the others in cancer progression. If we use this information to constrain the feature space and guide feature selection, we could potentially obtain more robust biomarkers. State-of-the-art studies have not utilized this knowledge but considered the whole feature space and the entire biological network. Because both the data and the network are large, the irrelevant information may overwhelm the signals. Furthermore, biological networks were typically integrated with one type of omics data. It would be very interesting to investigate how the prediction performance differs when the networks are integrated with different omics data types, and additionally what are the relationships among the features selected from different omics data.

To address this issue, we proposed a phenotype relevant network-based feature selection (PRNFS) framework. It consists of constructing a phenotype relevant gene regulatory network (GRN) and selecting features from this network. We demonstrated the superiority of this framework with the application of lung adenocarcinoma (LUAD) prognosis prediction. We constructed a GRN for EMT, which has been demonstrated as highly relevant to cancer metastasis and prognosis. On this network 4 types of omics data (mRNA-Seq, miRNA-Seq, DNA methylation, and copy number alteration data) were integrated and 10 feature selection algorithms were employed. We obtained both single- and multi-omics EMT prognostic signatures, evaluated their prediction performance, analyzed the biological interpretations, and performed survival analysis. Furthermore, these signatures were tested on independent multi-omics LUAD data. We showed that EMT prognostic signatures achieved remarkable prediction performance on TCGA data. On independent data, both single- and multi-omics signatures stratified patients into significantly different prognostic groups. Multi-omics signatures were shown to be more robust than single-omics signatures.

## Materials and methods

We will first describe the construction of EMT networks. This is followed by the introduction of 10 feature selection algorithms. Then we explain the details of the experiments.

### EMT gene regulatory networks

As an up-to-date EMT GRN is not readily available, we constructed the network by literature review. The network we constructed has incorporated key transcription factors, miRNAs, their regulations and interactions with EMT hallmark molecules. Multiple levels of gene regulations such as transcriptional, translational, and post-translational regulations were covered. The reference for each component in the network can be found in [37]. Since this network covers mainly driver genes, we named it as the *core network*.

As it is observed, driver genes are often less differentially expressed than the genes they regulate [35]. If one includes only the driver genes for identifying molecular signatures, one may have captured only partial information. We therefore extended this network by including the molecules that directly interact with or being regulated by the molecules in the core network. *NetworkAnalyst* tool [38] was employed to find these interactions, which consist of protein-protein interactions, miRNA-gene interactions, and transcription factor-gene interactions. The resulting network was named *extended network*. After constructing this network, we noticed that many features have a rather low variance among samples, we thus removed these features and obtained the *filtered network*. All three networks were employed in our experiments. The three networks contain 74, 123 and 455 nodes respectively. Details of the networks can be found in [37].

### Experiments

We first obtained RNA-Seq, miRNA-Seq, DNA methylation, and CNA data of LUAD from FIREHOSE Broad Genome Data Analysis Center website. mRNA-Seq and miRNA-Seq data were combined because they both measure the abundance of transcripts. This resulted in 3 data levels: gene expression, DNA methylation, and CNA data. These three data levels will be abbreviated as GE, DM, and CNA in the remaining text. Each data level was normalized feature-wise by subtracting the mean and dividing by the standard deviation. More details of data pre-processing can be found in [37, 39].

Since we have obtained 3 EMT networks and 3 data levels, feature selection can be performed on each combination of network and data level. To evaluate whether EMT-based feature selection can give more robust molecular signatures for prognosis prediction, we employed 10 representative features selection algorithms to identify signatures from EMT genes and EMT networks. Table 2 gives an overview of these algorithms. Five of these algorithms integrate network information and the other five algorithms use only omics data. The underlying methodologies are very different. We suppose that if EMT network is superior for selecting prognostic signatures, the performance of the selected features from the majority of these algorithms should show improvements. As mentioned before, state-of-the-art studies usually use only gene expression data for feature selection. We instead incorporated three different omics data levels. This gives us the possibility to compare and integrate the signatures from different data levels.

**Table 2.** Overview of the 10 feature selection algorithms. We listed below their main methodologies, whether network information was integrated, the algorithmic output and reference.

| Algorithm | Methodology | Network | Output | Reference |
| --- | --- | --- | --- | --- |
| t-test | t-statistic | No | feature ranking | [28, 40] |
| Lasso | regularized regression | No | coefficients | [41] |
| NetLasso | network-based regularization | Yes | coefficients | [15] |
| AddDA2 | subnetwork scoring and searching | Yes | subnetworks | [8, 12] |
| NetRank | feature importance on network | Yes | feature ranking | [19] |
| stSVM | random walk on network | Yes | feature ranking | [20] |
| Cox | Cox PH model | No | feature ranking | [42, 43] |
| RegCox | regularized Cox PH model | No | coefficients | [44] |
| MSS | random sampling | No | feature ranking | [45] |
| Survnet | subnetwork scoring and searching | Yes | subnetworks | [9] |

Note that even the largest EMT network (the extended network) covers only 2.3% of the original data dimensions. To assess the performance of EMT-based feature selection, we compared the prediction performance of EMT signatures with the features selected out of all features from the corresponding data levels. Additionally, random networks of the same size and structure as EMT networks were generated - the nodes in the random networks were randomly chosen from all the features of the corresponding data level.

The features selected from these random networks were compared with EMT signatures. In total, we compared the prediction performance of EMT signatures with the following 4 groups:

1. Random features. These features were selected from random networks using the same feature selection algorithms. 150 random networks were generated for feature selection.
2. All EMT features. We included all features in the EMT networks without feature selection.
3. All features from the corresponding data levels. This corresponds to 19,290 GE features, 20,074 DM features, or 21,456 CNA features.
4. Features selected from all data level features by applying Lasso algorithm.

We performed the comparison by selecting features with the training set, using these features to train an SVM classifier, and classifying samples on the cross-validation set. Patients who survived more than 1400 days belong to the good prognosis group and patients who survived less than 700 days belong to the poor prognosis group. The results from 30 times stratified 10-fold cross-validation were averaged. Within each data level the same cross-validation folds were used for all the feature selection algorithms on all the comparative groups. The classification performance was evaluated using three metrics: ROC-AUC, ROC-PR and accuracy.

We chose relatively stringent thresholds for feature selection, this is to reveal more difference than similarities between the two patient groups. We argue that it becomes harder to find the signatures if the two groups have more similar samples in terms of the phenotype. For example, if one uses a single threshold of 3 years, we assume that the molecular profiles of patients who survived a bit longer than 3 years may be very similar to patients who survived a bit shorter than 3 years. In this case, it is challenging to find the signatures that can capture the most important difference between the two groups, given the limited amount of samples and their heterogeneity. However, we did not omit the influence of thresholds. We tested the performance of all feature selection algorithms with four different thresholds, in the order of increasing discrepancy: 3 years, *<*900 or *>*1200 days, *<*700 or *>*1400 days, *<*500 or *>*1500 days.

Besides evaluating the classification performance, survival analysis was performed using the selected features on censored data. The data have much more samples that could not be included in classification.We think that if the selected features are good signatures, they should be able to stratify the patients into significantly different survival groups. We performed survival analysis on both all-stage and early-stage patients. The sample sizes for classification and for survival analysis (all stage patients) are given in Table 3.

**Table 3.** The description of datasets. This table shows the sample sizes for labeled data (thresholds *<*700 and *>*1400 days) and censored data.

|  | labeled data |  |  | censored data |
| --- | --- | --- | --- | --- |
|  | good prognosis | poor prognosis | total | all stages |
| Level GE | 84 | 99 | 183 | 497 |
| Level DM | 74 | 93 | 167 | 447 |
| Level CNA | 73 | 76 | 149 | 503 |

Last but not least, we analyzed the biological interpretations of EMT signatures. Instead of performing gene set enrichment analysis, which could give very significant results due to the biological context of the EMT networks, we employed association rule mining approach to infer prognostic association rules. The rules have the advantage to directly associate the states of the features to the phenotype of interest. We inferred rules using EMT signatures from individual data levels and also from their different combinations. Our motivation is to understand whether features from different data levels complement each other and jointly contribute to patient prognosis.

We were able to show that EMT signatures from different data levels complement each other in prognostic rules. This inspired us to obtain multi-omics EMT signatures by combining the signatures on individual data levels (single-omics signatures). Both single- and multi-omics EMT signatures were evaluated on TCGA data and independent LUAD multi-omics data using survival analysis. All the data and code for analysis are available at https://github.com/BorongShao/EMT prognosis-master.

## Results

### EMT signatures outperformed comparative groups

First, we show that regardless of the employed feature selection algorithms and evaluation metrics, EMT-based feature selection always outperforms feature selection on random networks. Fig 1 shows the distributions of AUC, AUPR, and accuracy values of EMT signatures and random ones, where DM data and core EMT network were used. S1 Fig shows the same comparative groups using GE data with filtered EMT network. In both cases, the advantages of EMT signatures are very apparent.

**Fig 1.**
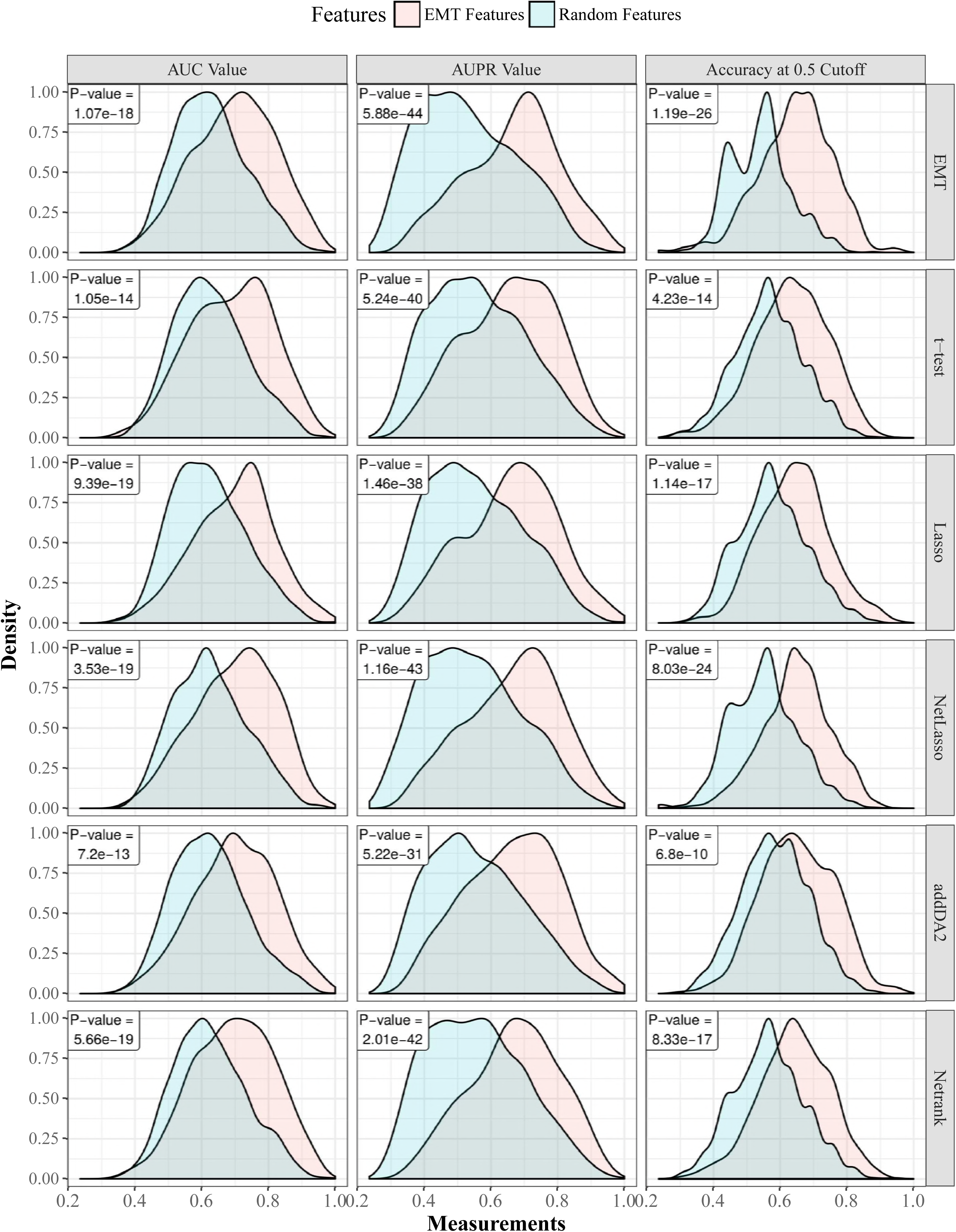
The AUC, AUPR, and accuracies of EMT features versus random features using DM data with the core EMT network. Gaussian kernel is used to estimate the density functions based on results from 30 times 10-fold cross-validation. For each cross-validation fold, EMT features and random features are tested on the same training and cross-validation samples. Each row in the figure corresponds to one feature selection algorithm. The last row corresponds to using all EMT features. The p-values of paired t-tests are provided in each sub-figure.

Next, we give the average AUC values of EMT signatures on all three data levels in Table 4. The boxplot is given in S2 Fig. These results show that features selected from GE and DM data obtained better prediction performance than features selected from CNA data. Depending on the data levels and network sizes, we find it hard to identify the best-performing feature selection algorithm. In the last two lines of the table we give the results of comparative groups 3 and 4. This shows that EMT signatures in many cases outperformed features selected from all data level features. For example, with Lasso feature selection algorithm, which was applied in both EMT feature space and in the whole feature space, EMT signatures gave better predictions in more than half of the cases. This indicates that selecting prognostic signatures from a much smaller phenotype relevant network is a feasible approach. We also evaluated the performance of the 10 feature selection algorithms with different classification thresholds. Both SVM and random forest classifiers are employed. The results are given in S3 Fig. It shows that regardless of feature selection algorithms, using more discrepant thresholds tends to obtain higher AUC values. Meanwhile, a few algorithms such as addDA2, RegCox, and Survnet are more sensitive to the effect of thresholds than the other algorithms.

**Table 4.** The prediction performance of EMT signatures on three data levels. The table gives the average AUC values of EMT signatures on three data levels with each EMT network. The results from comparative groups 2, 3, and 4 are given in the third row and the last two rows.

| Data Level | Gene expression |  |  | DNA Methylation |  |  | CNA |  |  |
| --- | --- | --- | --- | --- | --- | --- | --- | --- | --- |
| $\vert V(G)\vert $ | 74 | 123 | 455 | 74 | 123 | 455 | 70 | 117 | 445 |
| EMT | 0.662 | 0.728 | 0.691 | 0.698 | 0.679 | 0.671 | 0.616 | 0.645 | 0.608 |
| t-test | 0.658 | 0.709 | 0.677 | 0.688 | 0.675 | 0.669 | 0.616 | 0.626 | 0.621 |
| Lasso | 0.616 | 0.703 | 0.620 | 0.697 | 0.666 | 0.667 | 0.615 | 0.619 | 0.617 |
| NetLasso | 0.659 | 0.718 | 0.686 | 0.700 | 0.678 | 0.677 | 0.619 | 0.635 | 0.621 |
| addDA2 | 0.650 | 0.675 | 0.651 | 0.699 | 0.661 | 0.702 | 0.597 | 0.626 | 0.616 |
| Netrank | 0.656 | 0.691 | 0.668 | 0.695 | 0.685 | 0.693 | 0.615 | 0.619 | 0.610 |
| stSVM | 0.651 | 0.693 | 0.639 | 0.669 | 0.668 | 0.687 | 0.608 | 0.617 | 0.616 |
| Cox | 0.673 | 0.705 | 0.712 | 0.703 | 0.707 | 0.696 | 0.620 | 0.664 | 0.675 |
| RegCox | 0.648 | 0.698 | 0.729 | 0.696 | 0.717 | 0.666 | 0.645 | 0.669 | 0.653 |
| MSS | 0.662 | 0.694 | 0.659 | 0.674 | 0.654 | 0.640 | 0.608 | 0.627 | 0.625 |
| Survnet | 0.646 | 0.661 | 0.679 | 0.702 | 0.688 | 0.680 | 0.626 | 0.693 | 0.682 |
| All | 0.648 |  |  | 0.652 |  |  | 0.612 |  |  |
| All + Lasso | 0.643 |  |  | 0.691 |  |  | 0.607 |  |  |

### Frequently selected features further improves predictions

Although EMT signatures were shown to be significantly predictive in the experiments above, we observed high variance in the AUC values from individual cross-validation tests (shown in S2 Fig). Some partitions of data into training and cross-validation sets led to good predictions and some led to poor predictions. Even on the small EMT feature space, this phenomenon is already frequently observed. This suggests that selecting molecular signatures based on single cross-validation test or single sample division into training and testing set is highly unreliable.

We think that sample heterogeneity contributed to the high variance in prediction performance. Thus, we addressed this issue by employing the frequently selected features (FSFs) from all 30 times 10-fold cross-validation feature selection. Instead of using 20 features selected from each training set, we used the top 20 FSFs and tested their performance using the same evaluation approach. DM data and the extended EMT network were employed for the test, as this combination was shown in Table 4 and Figure 4 to give above-average prediction performance. We compared the prediction performance of FSFs with that of individually selected features. The results are given in Fig 2. The density plots and results of statistical tests are given in S4 Fig.

**Fig 2.**
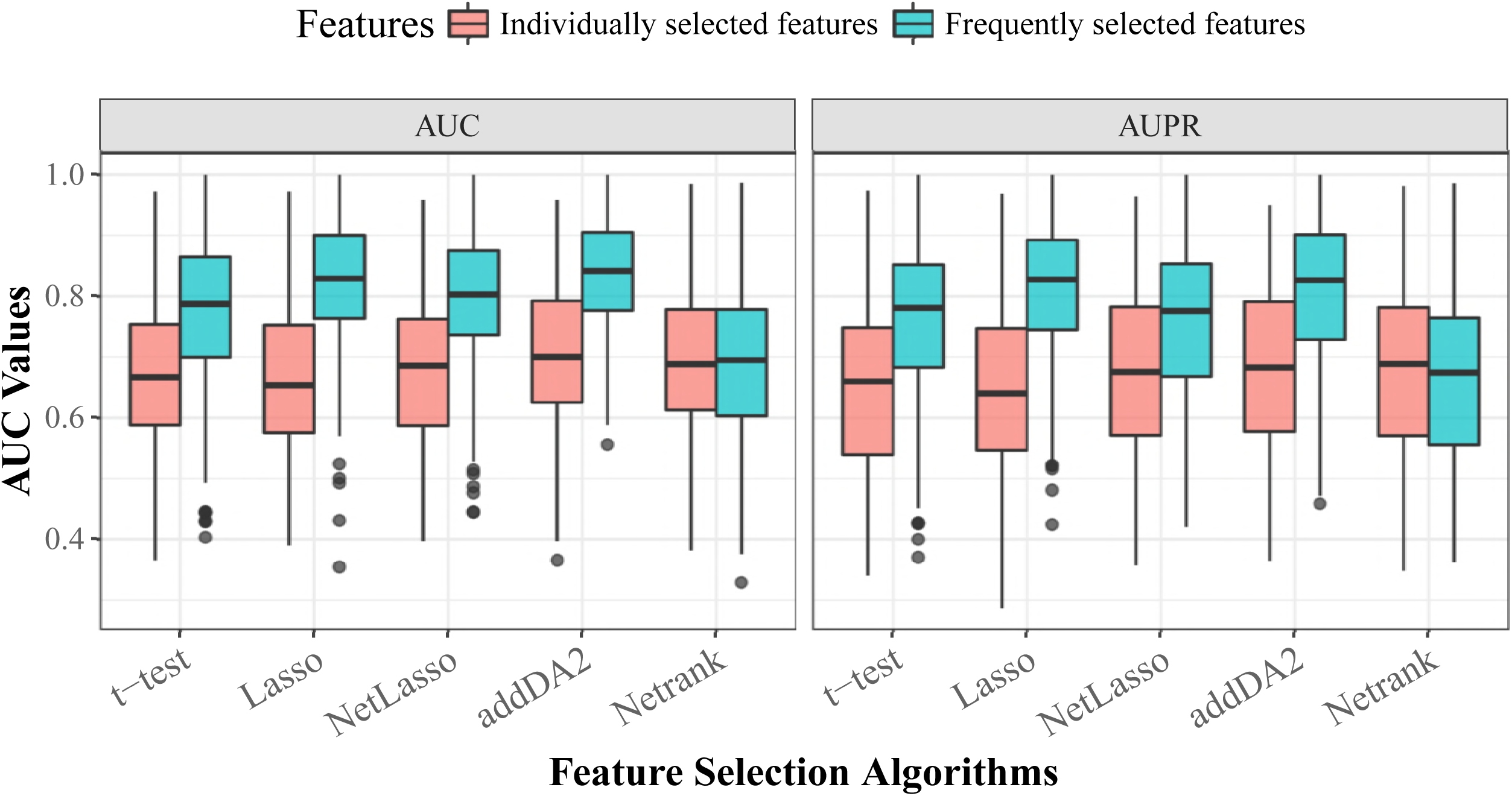
The comparison of prediction performance between FSFs and individually selected features for different feature selection algorithms. The boxplot is based on the results from 30 times stratified 10-fold cross-validation.

**Fig 3.**
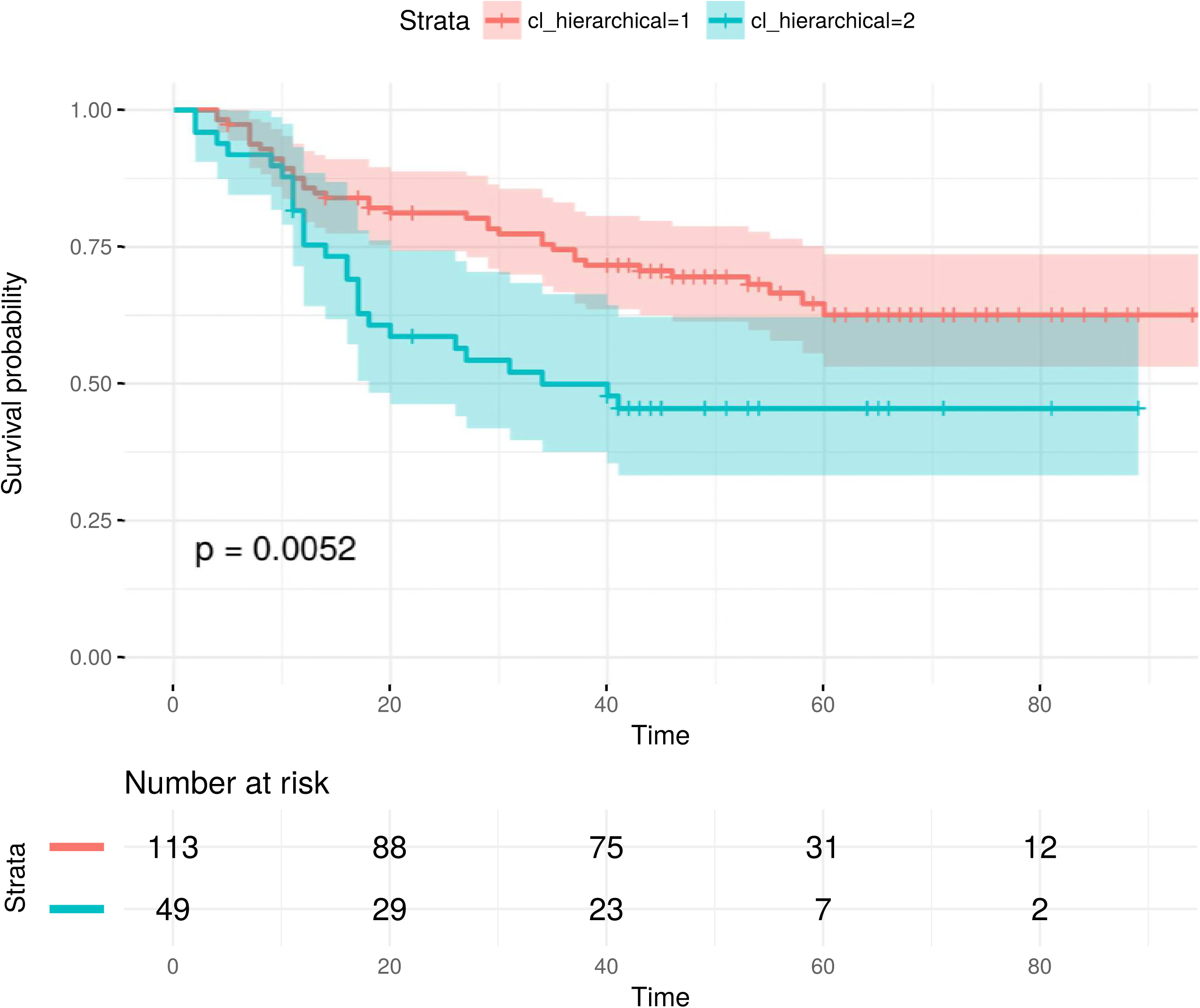
EMT single-omics signatures can stratify test samples into significantly different prognostic groups. The signature is selected by addDA2 algorithm using DM data.

**Fig 4.**
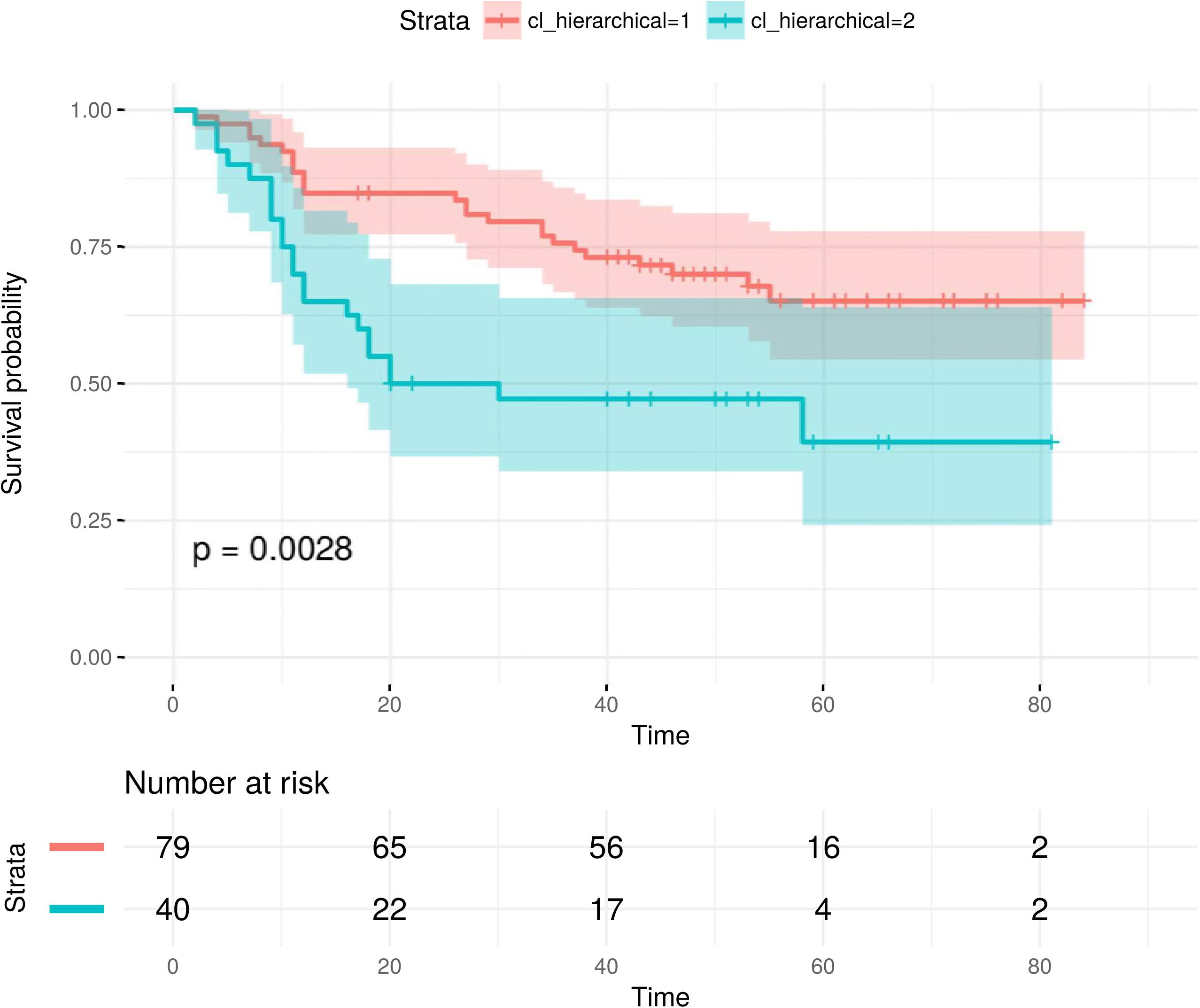
EMT multi-omics signatures can stratify test samples into significantly different prognostic groups, when the corresponding single-omics signatures cannot. The signature consists of both GE and DM single-omics signatures selected by t-test.

We observed that FSFs significantly outperformed individually selected features, except for Netrank algorithm. The average AUC values of t-test, Lasso, NetLasso, and addDA2 feature selection algorithms were 0.773, 0.825, 0.796, and 0.833, respectively. It shows that using FSFs can mitigate the effect of sample heterogeneity. Recall that we used only *<*2.5% of the original dimensionality, namely EMT features, for feature selection and prognosis prediction. The remarkable results are consistent with biological knowledge that EMT process is highly relevant to cancer prognosis [46–50].

### Biological interpretations

After identifying EMT FSFs, we further investigated their biological interpretations, especially the relationships among FSFs from different omics data levels. We employed association rule mining approach [51]. It is originally defined as the following [52]:

Let *I* = *{i*_1_, *i*_2_, *…, i*_*n*_*}* be a set of *n* binary features called items. Let *D* = *{t*_1_, *t*_2_, *…, t*_*m*_*}* be a set of transactions called the database. A rule is defined in the form: *X ⇒Y*, where *X ⇒ Y, ⊆ I*. The itemsets *X* and *Y* are called left-hand-side (LHS) and right-hand-side (RHS). In order to select interesting rules from the set of all possible rules, constraints on various measures of significance and interest are applied. Let a rule *X ⇒ Y* be identified on a set of transactions *T*. Commonly used constraints are given below:

- Support. It indicates how frequently the itemset appears in *T*.

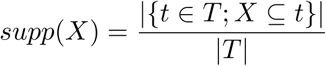
- Confidence. It indicates how often a rule has been found to be true.

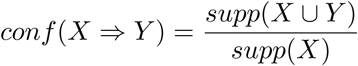
- Lift. It indicates the degree to which *X* and *Y* depend on each other.

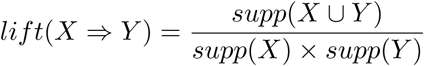

We binarized the EMT features using their means and applied Apriori algorithm [53] to derive rules, with the constraints of *confidence ≥* 0.8 and *support* ≥ 0.1. The algorithm was implemented in the textitarules R package [51]. Since we are trying to find molecular patterns for predicting prognosis, we set the RHS of the rules to be the class labels of prognosis.

The resulting rules show sound biological interpretations according to established findings in cancer research. Here we interpret two rules identified from the core EMT network:

*{LOXL*2_*GE*_ = *high, TGFB*1_*GE*_ = *high, miR −* 34*a*_*GE*_ = *low} ⇒ {prognosis* = *poor}*, with *support* = 0.135, *confidence* = 1, *lift* = 2.046. This rule applies to all samples that have these 3 gene expression conditions. Biologically, it has been shown that LOXL2 can stabilize SNAI1. TGFB1 can phosphorylate SMAD2 and SMAD3, which interact with SMAD4 to activate HMGA2, which then activates SNAI1. When LOXL2 and TGFB1 are highly expressed, it not only induces SNAI1 gene expression but also stabilizes SNAI1 protein. miR-34a has the role of repressing SNAI1. When miR-34a has low gene expression, SNAI1 is less repressed. Taken together, these three conditions point to the direction of the high expression of SNAI1 - a master transcription factor to induce EMT. This contributes to poor prognosis. In contrast, another rule which has an opposite LOXL2 state indicates good prognosis:

*{LOXL*2_*GE*_ = *low, ETS*1_*GE*_ = *low, LOXL*2*_DM_* = *high}⇒ prognosis* = *good*, with *support* = 0.105, *confidence* = 1, *lift* = 1.956. In this scenario, LOXL2 has high DNA methylation level and low gene expression level, and thus not able to stabilize SNAI1. ETS1 gene is known to increase the expression of ZEB1 which induces EMT. In this rule ETS1 has low expression so it does not contribute to inducing EMT. These factors can contribute to good prognosis. S1 Table contains more examples.

### From single- to multi-omics signatures

The FSFs above were obtained alternatively from single data levels. Therefore, we name them as *single-omics* signatures. To investigate whether molecular signatures incorporating multiple data levels can be superior, we combined single-omics signatures into *multi-omics* signatures and compared their capabilities in stratifying samples into different prognostic groups. Using these signatures, we clustered the samples into 3 groups with both k-means and spectral clustering algorithms. Survival analysis was performed on the resulting cluster by estimating Kaplan-Meier survival curves and conducting log-rank tests. The test results based on k-means algorithm are given in Table 5, where the columns show different data level combinations and the rows correspond to feature selection algorithms. The comparative group of using all EMT features without feature selection is included. The test results based on spectral clustering algorithm are given in S2 Table. In both tables we observe that multi-omics signatures improve sample stratifications significantly. An example is visualized in S5 Fig, S6 Fig, and S7 Fig.

**Table 5.** The results of log-rank tests on stratified sample clusters using single- and multi-omics EMT signatures on all-stage samples. K-means algorithm was employed for clustering the samples into 3 groups. We highlighted all p-values that are lower than 10e-3.

|  | GE | DM | CNA | GE+DM | GE+CNA | DM+CNA | GE+DM+CNA |
| --- | --- | --- | --- | --- | --- | --- | --- |
| t-test | **7.87e-06** | 1.12e-01 | 3.62e-03 | **7.55e-06** | **8.75e-04** | 1.90e-03 | **8.25e-06** |
| Lasso | **4.58e-06** | 5.56e-02 | 5.54e-01 | **1.66e-04** | **2.28e-07** | 8.71e-01 | **1.18e-04** |
| NetLasso | 2.71e-01 | 6.45e-01 | 7.58e-02 | 2.96e-02 | 1.53e-02 | 4.83e-01 | 4.34e-01 |
| addDA2 | **5.20e-10** | **5.17e-09** | **1.24e-04** | **8.99e-18** | **3.75e-05** | **1.11e-09** | **1.35e-07** |
| Netrank | **1.13e-07** | 1.79e-01 | 2.19e-02 | **2.50e-07** | **6.97e-06** | 7.31e-02 | **3.28e-06** |
| stSVM | 4.14e-02 | 3.39e-01 | 8.91e-01 | 8.86e-01 | 6.37e-02 | 5.85e-01 | 6.83e-01 |
| Cox | **2.55e-09** | 2.91e-03 | **5.30e-07** | **3.12e-04** | **1.70e-06** | **1.13e-04** | **6.11e-06** |
| RegCox | **1.78e-07** | 8.52e-03 | 2.67e-01 | **2.36e-09** | **1.52e-10** | **1.81e-07** | **2.52e-07** |
| MSS | 1.48e-03 | 5.95e-01 | 2.78e-01 | **6.29e-05** | **2.28e-04** | 2.59e-01 | 1.63e-03 |
| Survnet | **7.36e-05** | 6.25e-03 | 5.19e-03 | **2.59e-06** | 1.54e-03 | **3.77e-05** | **2.19e-05** |
| Ensemble | **2.32e-09** | 4.72e-02 | 6.05e-03 | **1.20e-04** | **1.62e-05** | 1.01e-01 | **1.18e-04** |
| allemt | 1.39e-02 | 4.30e-01 | 1.07e-01 | 7.64e-01 | 1.45e-02 | 2.07e-01 | 5.34e-01 |

Next, we performed survival analysis on early stage patients. The results are given in Table 6. It shows that EMT-based signatures can still stratify the patients into significantly different prognostic groups.

**Table 6.**
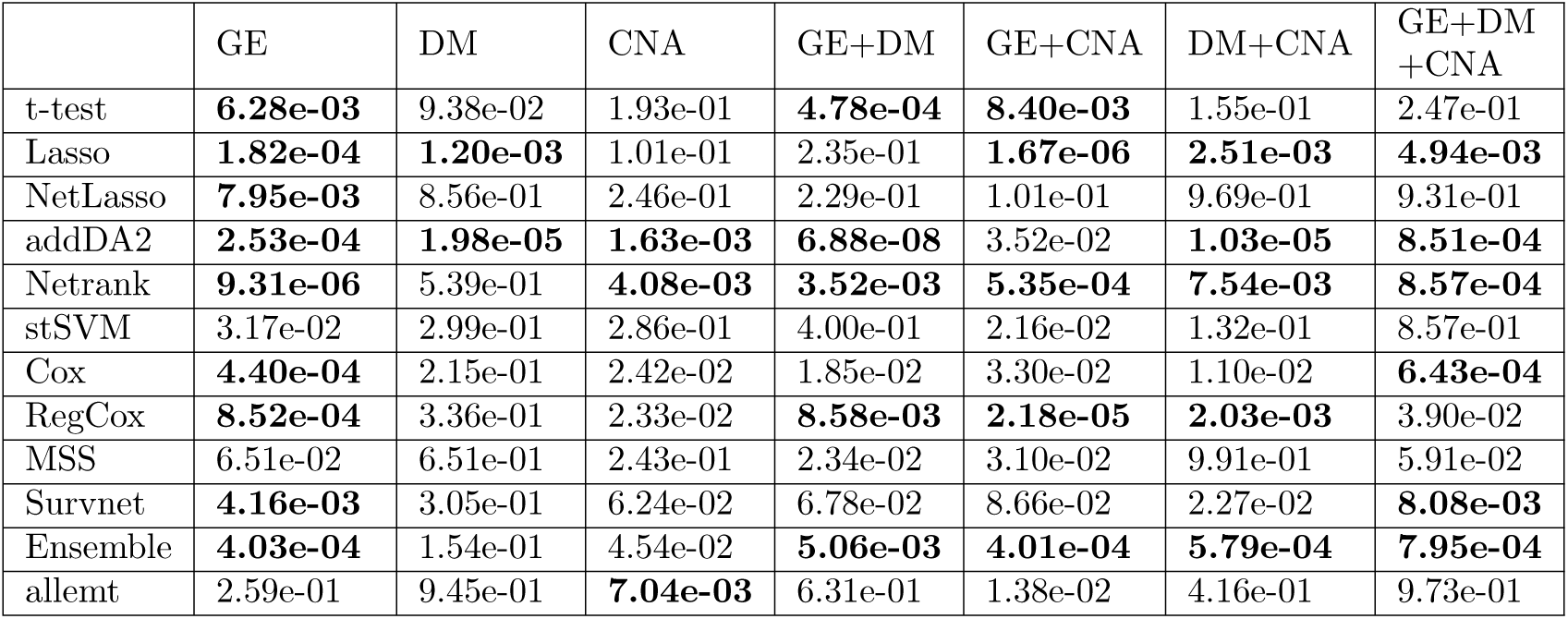
The results of log-rank tests on stratified sample clusters using single- and multi-omics EMT signatures on early-stage samples. K-means algorithm was employed for clustering the samples into 3 groups. We highlighted all p-values that are lower than 10e-2.

|  | GE | DM | CNA | GE+DM | GE+CNA | DM+CNA | GE+DM+CNA |
| --- | --- | --- | --- | --- | --- | --- | --- |
| t-test | **6.28e-03** | 9.38e-02 | 1.93e-01 | **4.78e-04** | **8.40e-03** | 1.55e-01 | 2.47e-01 |
| Lasso | **1.82e-04** | **1.20e-03** | 1.01e-01 | 2.35e-01 | **1.67e-06** | **2.51e-03** | **4.94e-03** |
| NetLasso | **7.95e-03** | 8.56e-01 | 2.46e-01 | 2.29e-01 | 1.01e-01 | 9.69e-01 | 9.31e-01 |
| addDA2 | **2.53e-04** | **1.98e-05** | **1.63e-03** | **6.88e-08** | 3.52e-02 | **1.03e-05** | **8.51e-04** |
| Netrank | **9.31e-06** | 5.39e-01 | **4.08e-03** | **3.52e-03** | **5.35e-04** | **7.54e-03** | **8.57e-04** |
| stSVM | 3.17e-02 | 2.99e-01 | 2.86e-01 | 4.00e-01 | 2.16e-02 | 1.32e-01 | 8.57e-01 |
| Cox | **4.40e-04** | 2.15e-01 | 2.42e-02 | 1.85e-02 | 3.30e-02 | 1.10e-02 | **6.43e-04** |
| RegCox | **8.52e-04** | 3.36e-01 | 2.33e-02 | **8.58e-03** | **2.18e-05** | **2.03e-03** | 3.90e-02 |
| MSS | 6.51e-02 | 6.51e-01 | 2.43e-01 | 2.34e-02 | 3.10e-02 | 9.91e-01 | 5.91e-02 |
| Survnet | **4.16e-03** | 3.05e-01 | 6.24e-02 | 6.78e-02 | 8.66e-02 | 2.27e-02 | **8.08e-03** |
| Ensemble | **4.03e-04** | 1.54e-01 | 4.54e-02 | **5.06e-03** | **4.01e-04** | **5.79e-04** | **7.95e-04** |
| allemt | 2.59e-01 | 9.45e-01 | **7.04e-03** | 6.31e-01 | 1.38e-02 | 4.16e-01 | 9.73e-01 |

Last but not least, we tested the performance of two integrative clustering algorithms: SNF [54] and iCluster [55] with multi-omics EMT signatures. Briefly, SNF algorithm constructs sample similarity networks using individual data levels and then fuses these networks into a single similarity network, where spectral clustering is used to decide sample clusters. iCluster employs joint latent variable model to connect different data levels. The latent component is used to determine sample clusters. Based on the clustering results we performed survival analysis. The results of log-rank tests are given in S3 Table for SNF algorithm and in S4 Table for iCluster algorithm. We observed that neither SNF nor iCluster algorithm yielded better sample stratifications than using k-means algorithm (Table 5).

### Test results on independent data

We obtained the test data from [56] including 164 samples with DM data. 121 of these samples have also mRNA expression data (microarray) available. The patient follow up time ranges between 2 and 99 months with the median of 44 months. The outcome (event) is defined as the occurrence of relapse, distant metastasis or death. The time to event is calculated from the date of surgery. Detailed experimental procedures and the processing of raw data are provided in [56]. EMT single-and multi-omics signatures consisting of GE and DM data levels were evaluated on the test data using survival analysis. EMT signatures were extracted from the test data without any additional training or modifications. Hierarchical clustering, instead of k-means was employed in order to compare our results with the original study [56].

We have tested the EMT signatures selected by each feature selection algorithm [37]. It is show that single-omics signatures can already stratify the samples into significantly different prognostic groups. An example is given in Fig 3. Multi-omics signatures often yielded better sample stratifications. Fig 4 shows an example where the multi-omics signature from a feature selection algorithm can significantly stratify the samples while the single-omics signatures cannot. Compared with the survival analysis results in the original study [56], we achieved more significant sample stratifications with EMT signatures.

## Discussion

Various feature selection algorithms have been proposed to identify biomarkers from Omics data for predicting the phenotype of interest. Although more and more information such as biological networks and multiple types of omics data have been integrated in feature selection, recent studies show the low reproducibility of molecular signatures [22, 24, 35]. Some accredit this to the existence of a large number of genes that are correlated with the target labels [12]. Given the limited amount of samples, it becomes very hard to differentiate the marker genes and irrelevant genes. We addressed this issue by constructing a phenotype relevant gene regulatory network, integrating multiple types of omics data with the network to select molecular signatures. We have shown that with lung cancer prognosis prediction, EMT signatures selected from only 2.5% of the original feature space outperformed the classical feature selection on the whole feature space. To the best of our knowledge, we for the first time constructed a phenotype-relevant GRN for lung cancer prognosis prediction.

Previously we employed EMT networks for selecting lung cancer prognostic signatures [39, 57]. However, [57] used mRNA expression and miRNA expression data only. [39] employed three data levels for feature selection but obtained no significant improvement in predictions. In this study, we extended the EMT network to incorporate its interacting molecules. Besides, we reviewed the network used in [39] and removed the edges which denote associations rather than direct gene regulations. What also distinguishes this study from our previous work is the employment of 10 representative feature selection algorithms, instead of decomposing the network into network motifs [39, 57]. We have selected EMT signatures on three data levels with different network sizes, compared with the features selected from the whole data dimensions, and derived prognostic rules from EMT signatures. Furthermore, we obtained multi-omics signatures and showed their superior prediction performance over single-omics signatures. This shows that signatures from multiple omics data types can complement each other to better distinguish different phenotypes.

The potential of EMT molecules in prognosis prediction has also been studied before.[61] and [62] performed survival analysis using individual EMT hallmark molecules such as E-cadherin and vimentin and showed that none of these molecules could separate LUAD or bladder cancer patients into significantly different prognostic groups. Note that these conclusions were drawn from mainly univariate analysis. Since the molecules jointly contribute to the phenotype, it could be more helpful to use a set of features. This can be seen also from the prognostic association rules derived from EMT signatures, where EMT molecules are jointly associated with the phenotype. All in all, we successfully demonstrated that EMT network-based feature selection and data integration can provide advantages in selecting cancer prognostic signatures.

## Supporting information

**Fig S1.**
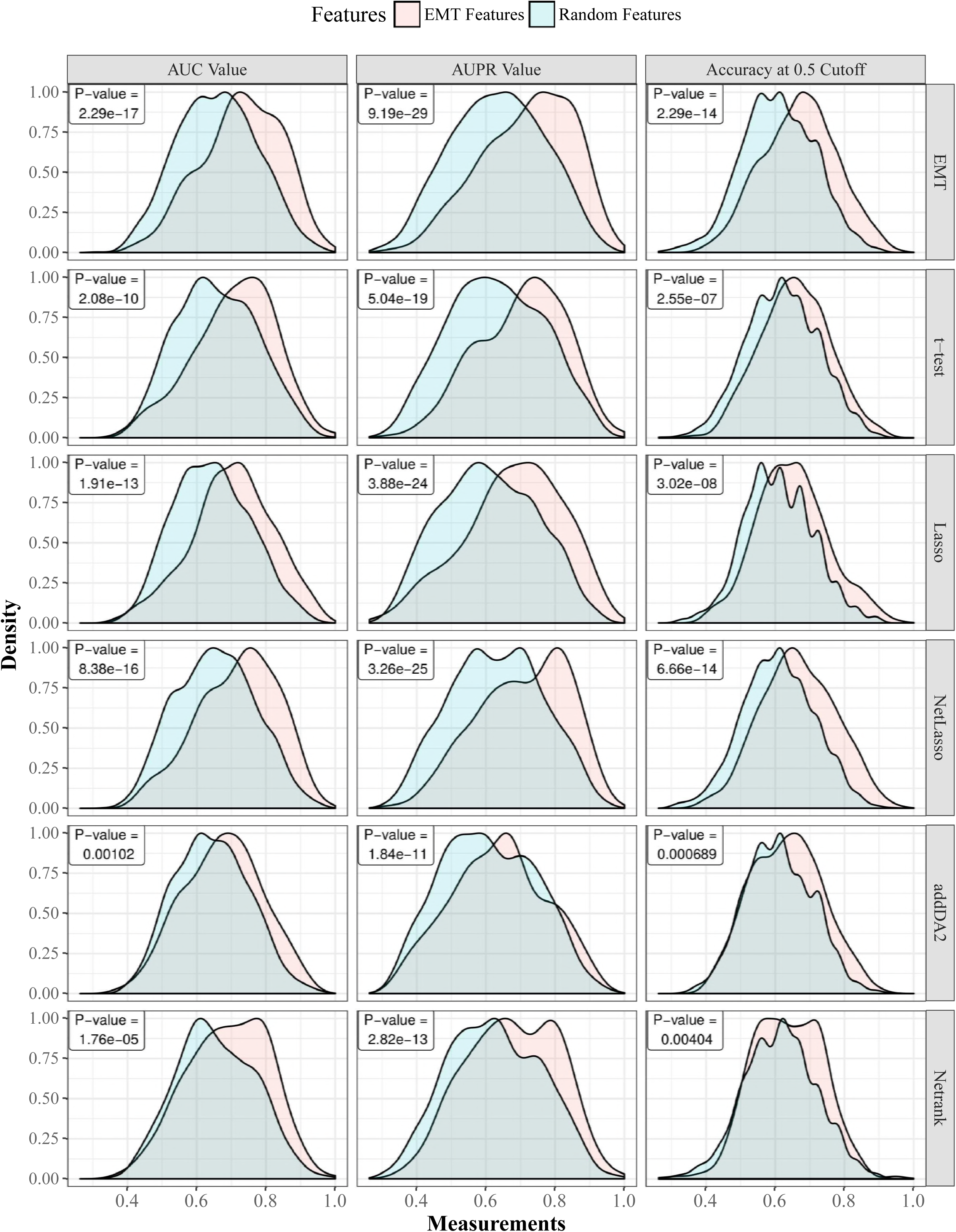
The AUC, AUPR, and accuracies of EMT features versus random features using gene expression data with filtered EMT network. Gaussian kernel is used to estimate the density functions based on results from 30 times 10-fold cross-validation. For each cross-validation fold, EMT features and random features are tested on the same training and testing samples. The comparisons on five feature selection algorithms together with the comparative group of using all EMT features are shown. The p-values of paired t-tests are provided.

**Fig S2.**
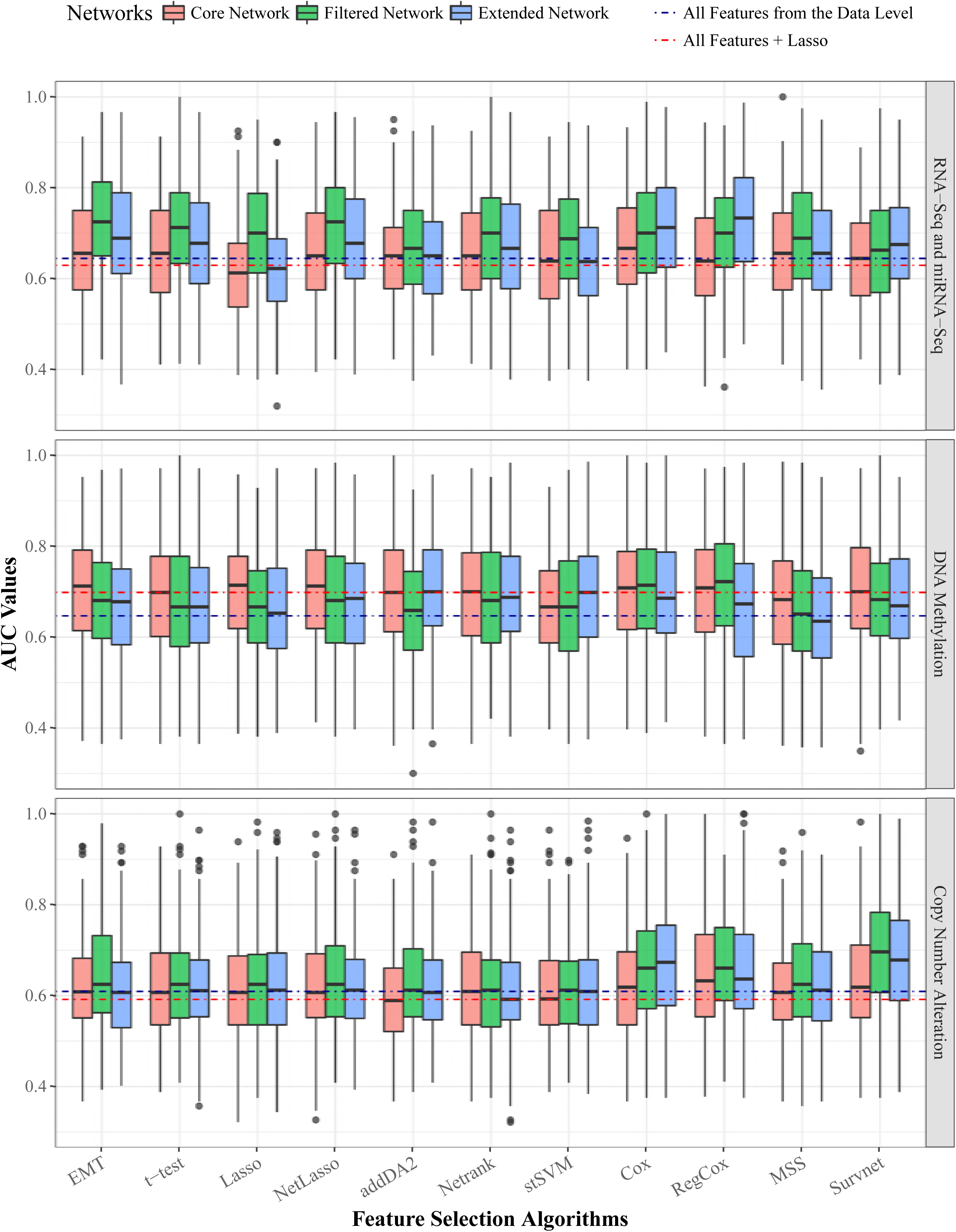
The AUC values of 10 feature selection algorithms. The three panels correspond to three data levels. Within each panel, the AUC values of the 10 algorithms are plotted. Each algorithm has three boxes of different colors denoting the 3 EMT networks. The blue and red dotted lines within each panel are the median AUC values of two comparative groups: 1) using all data level features and 2) Lasso feature selection on all data level features.

**Fig S3.**
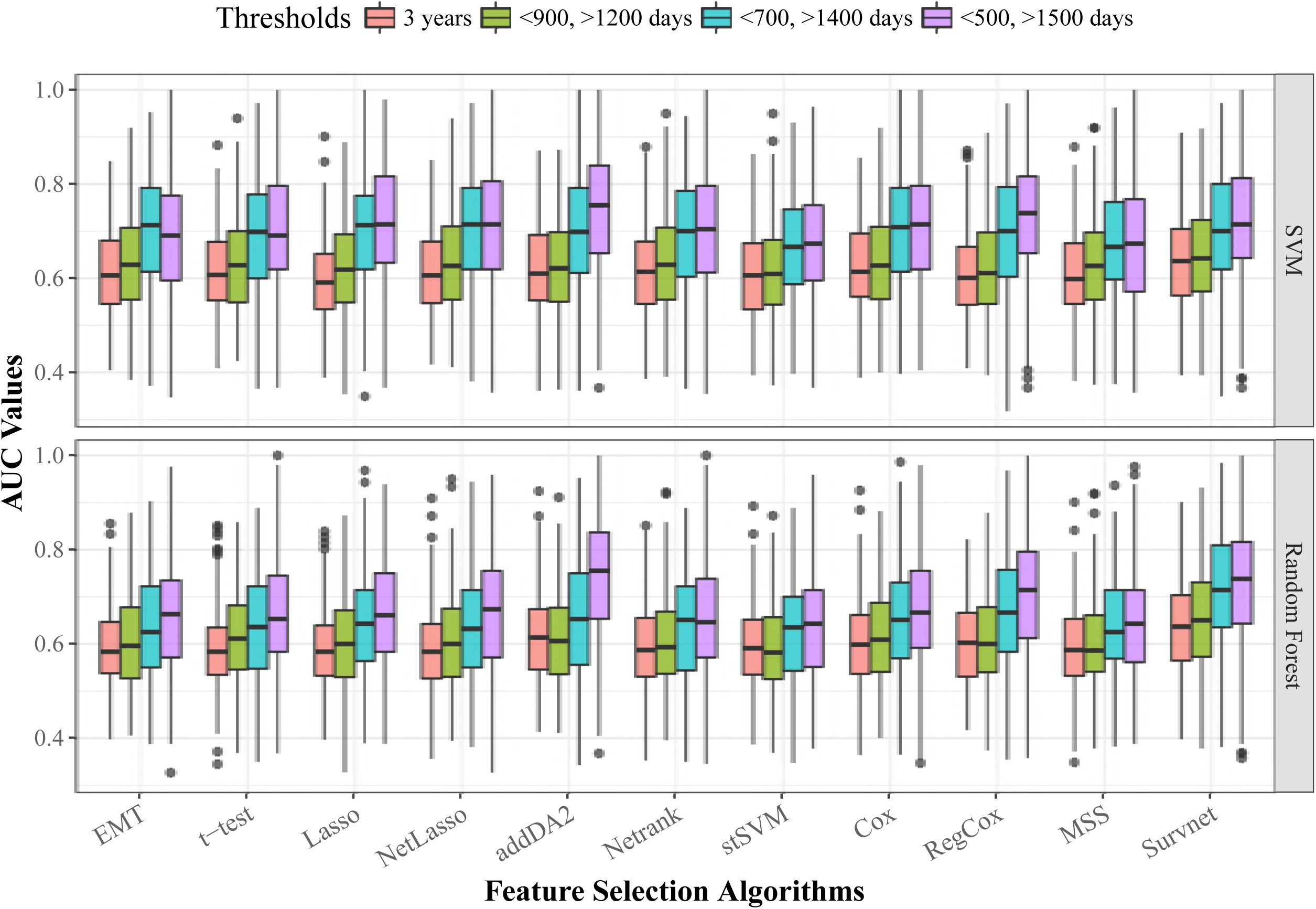
The AUC values of 10 feature selection algorithms using different thresholds for classification. The data level is DNA methylation data. The network is EMT core network.

**Fig S4.**
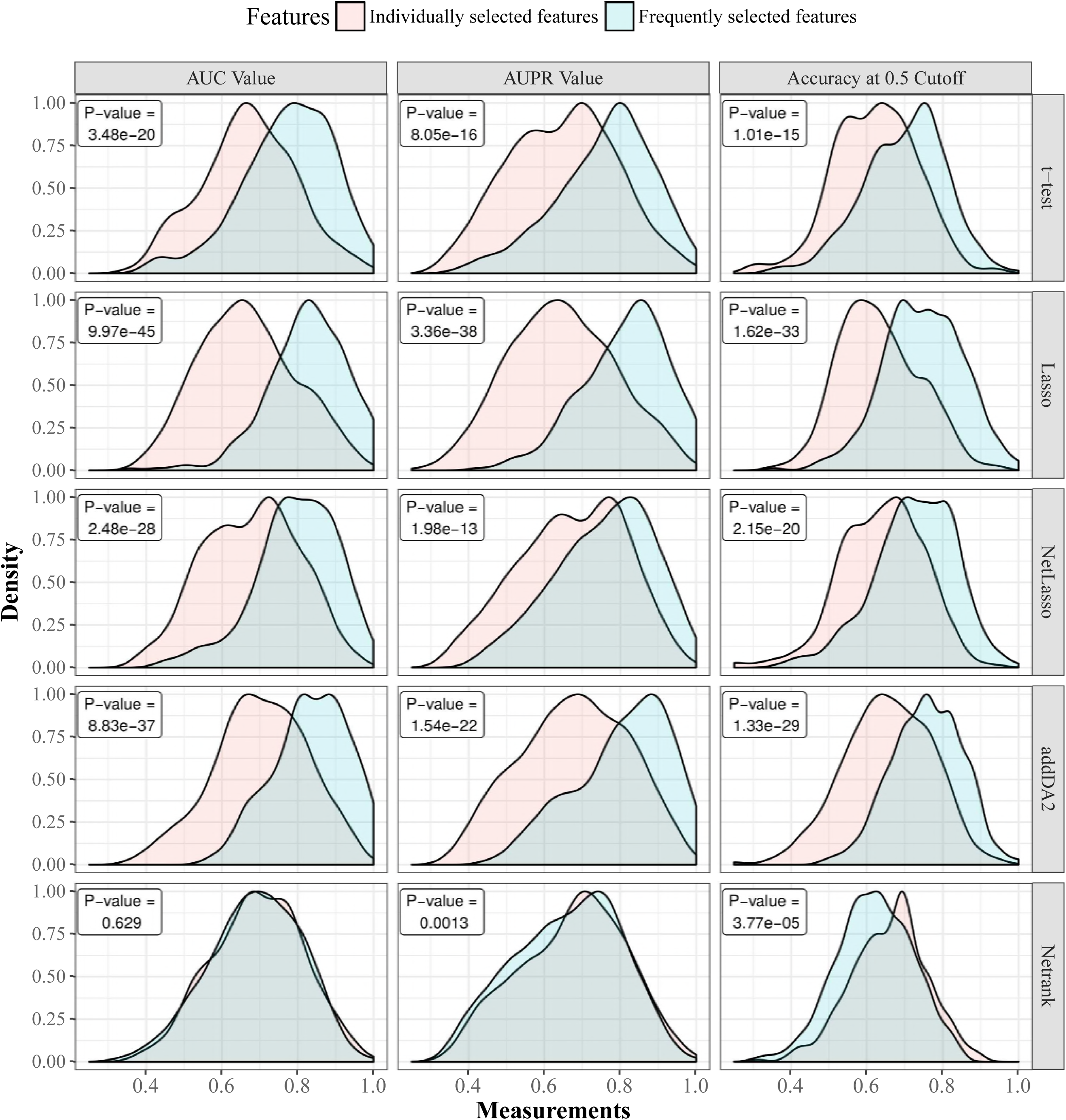
The comparison of FSFs with individually selected features in terms of AUC, AUPR, and accuracy values. We used DNA methylation data and extended EMT network for feature selection and SVM classifier for classification. Gaussian kernel is used to estimate the density functions based on results from 30 times stratified 10-fold cross-validation. For each cross-validation iteration, individually selected features and FSFs are tested on the same training and testing samples. The comparison between the two feature groups is shown on five feature selection algorithms together with the p-values of paired t-tests.

**Fig S5.**
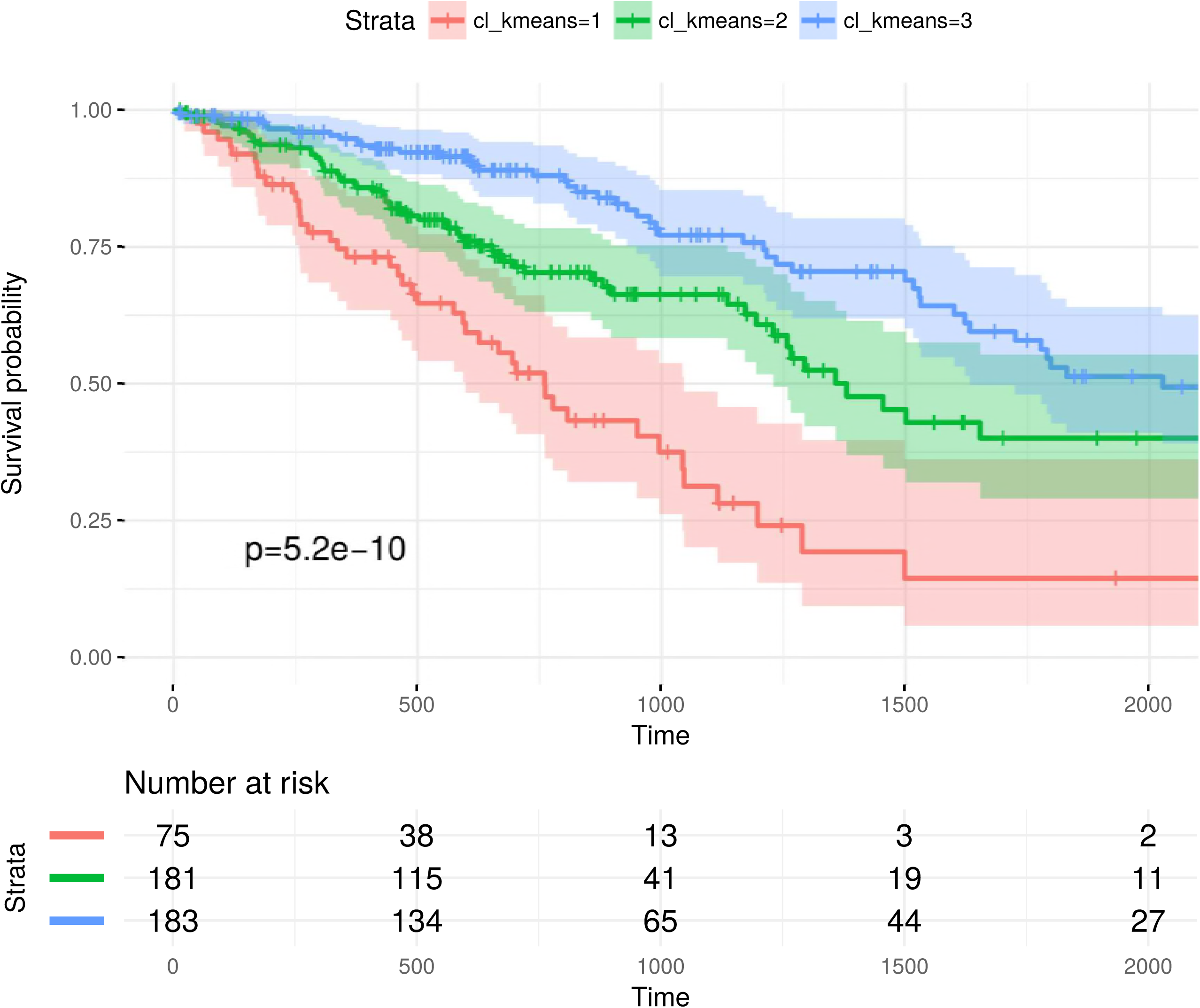
Patient stratification using GE features from addDA2 algorithm.

**Fig S6.**
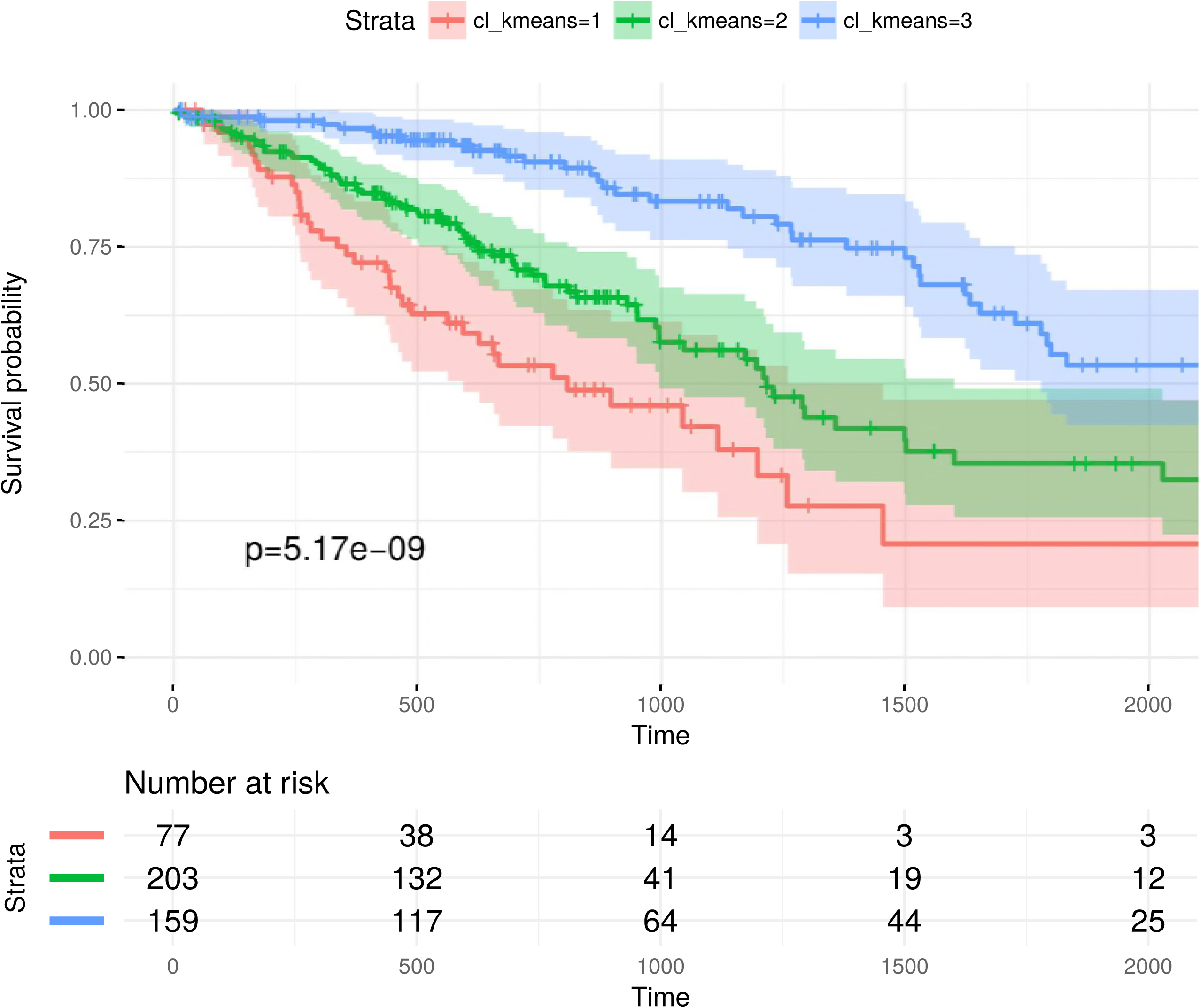
Patient stratification using DM features from addDA2 algorithm.

**Fig S7.**
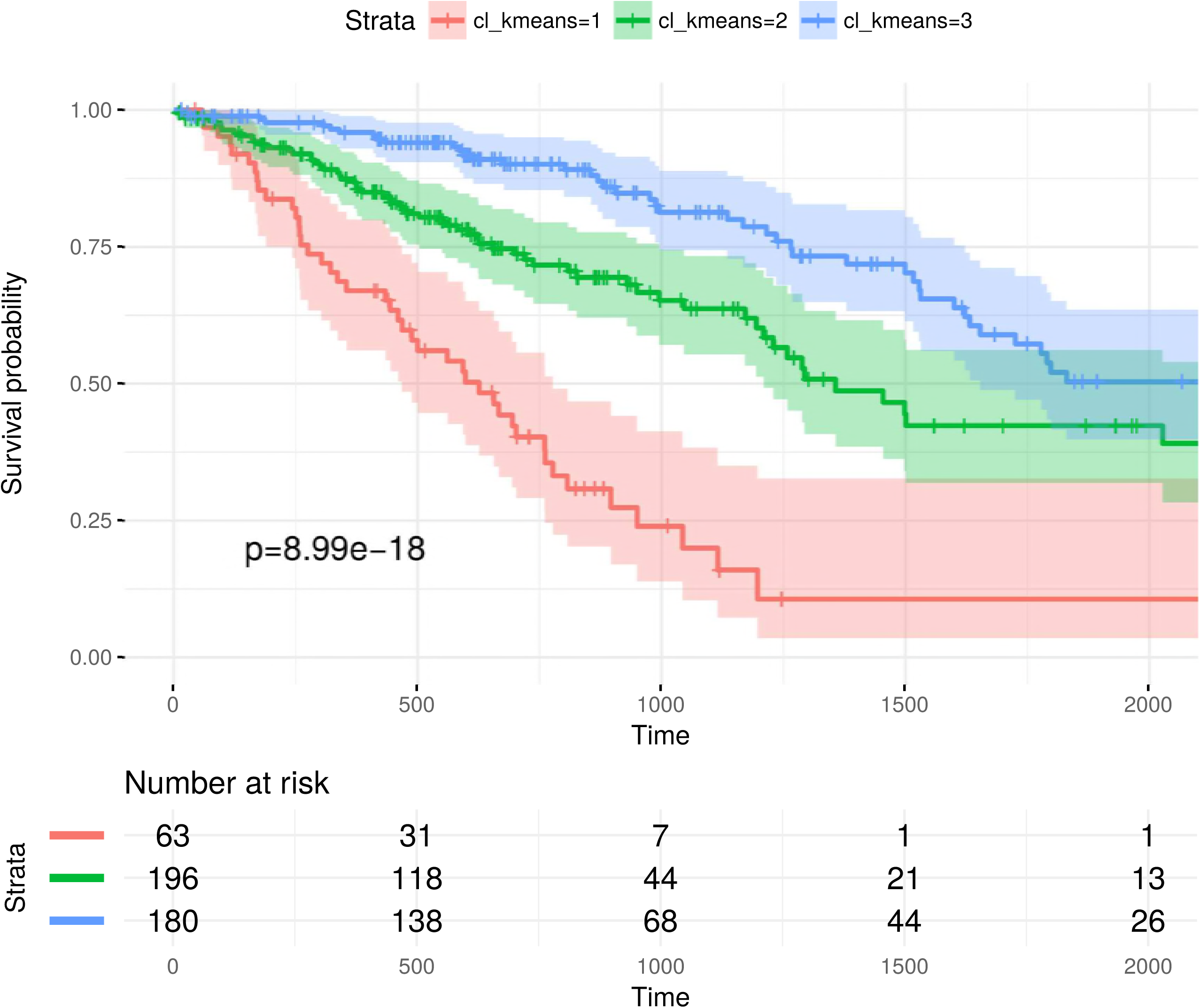
Patient stratification using GE and DM features from addDA2 algorithm.

**Table S1.** Top 20 prognostic association rules derived from the FSFs using filtered EMT network. All the following rules have confidence scores of 1.

|  | LHS | prognosis | supp | lift |
| --- | --- | --- | --- | --- |
| 1 | $CDH3_{GE} = low, miR.34a_{DM} = high, HMGA2_{DM} = low$ | good | 0.113 | 1.956 |
| 2 | $EGLN3_{GE} = low, CDH3_{GE} = low, HMGA2_{DM} = low$ | good | 0.113 | 1.956 |
| 3 | $GFI1B_{GE} = high, CDH3_{GE} = low, HMGA2_{DM} = low$ | good | 0.135 | 1.956 |
| 4 | $GATA6_{GE} = low, E2F1_{GE} = low, HMGA2_{DM} = low$ | good | 0.120 | 1.956 |
| 5 | $GATA6_{GE} = high, CDC20_{GE} = low, FOXA3_{GE} = high$ | poor | 0.150 | 2.046 |
| 6 | $GATA6_{GE} = high, CDH3_{GE} = high, FOXA3_{GE} = high$ | poor | 0.135 | 2.046 |
| 7 | $GATA6_{GE} = high, miR.34a_{DM} = low, FOXA3_{GE} = high$ | poor | 0.143 | 2.046 |
| 8 | $miR.34a_{GE} = low, CCND1_{GE} = high, FOXA3_{GE} = low$ | good | 0.113 | 1.956 |
| 9 | $GATA6_{GE} = low, miR.34a_{GE} = low, CCND1_{GE} = high$ | good | 0.105 | 1.956 |
| 10 | $BIRC3_{GE} = low, CCND1_{GE} = high, FOXA3_{GE} = low$ | good | 0.113 | 1.956 |
| 11 | $LOXL2_{GE} = high, miR.34a_{DM} = low, FOXA3_{GE} = high$ | poor | 0.158 | 2.046 |
| 12 | $BIRC3_{GE} = low, miR.34a_{GE} = low, CDH3_{GE} = low$<br>$HMGA2_{DM} = low$ | good | 0.105 | 1.956 |
| 13 | $GFI1B_{GE} = high, miR.34a_{GE} = low, HMGA2_{DM} = low$<br>$FOXA3_{GE} = low$ | good | 0.105 | 1.956 |
| 14 | $GFI1B_{GE} = high, BIRC3_{GE} = low, miR.34a_{GE} = low$<br>$HMGA2_{DM} = low$ | good | 0.105 | 1.956 |
| 15 | $ITGA6_{GE} = high, BIRC3_{GE} = high, BIRC5_{GE} = high$<br>$GATA6_{GE} = high$ | poor | 0.113 | 2.046 |
| 16 | $BIRC3_{GE} = high, GATA6_{GE} = high, E2F1_{GE} = low$<br>$miR.34a_{DM} = low$ | poor | 0.105 | 2.046 |
| 17 | $BIRC3_{GE} = high, GATA6_{GE} = high, E2F1_{GE} = low$<br>$GATA4_{GE} = low$ | poor | 0.135 | 2.046 |
| 18 | $BIRC5_{GE} = high, GATA6_{GE} = high, miR.34a_{GE} = high$<br>$FOXA3_{GE} = high$ | poor | 0.128 | 2.046 |
| 19 | $EGLN3_{GE} = high, BIRC5_{GE} = high, GATA6_{GE} = high$<br>$FOXA3_{GE} = high$ | poor | 0.113 | 2.046 |
| 20 | $BIRC5_{GE} = high, GATA6_{GE} = high, miR.192_{GE} = low$<br>$FOXA3_{GE} = high$ | poor | 0.120 | 2.046 |

**Table S2.**
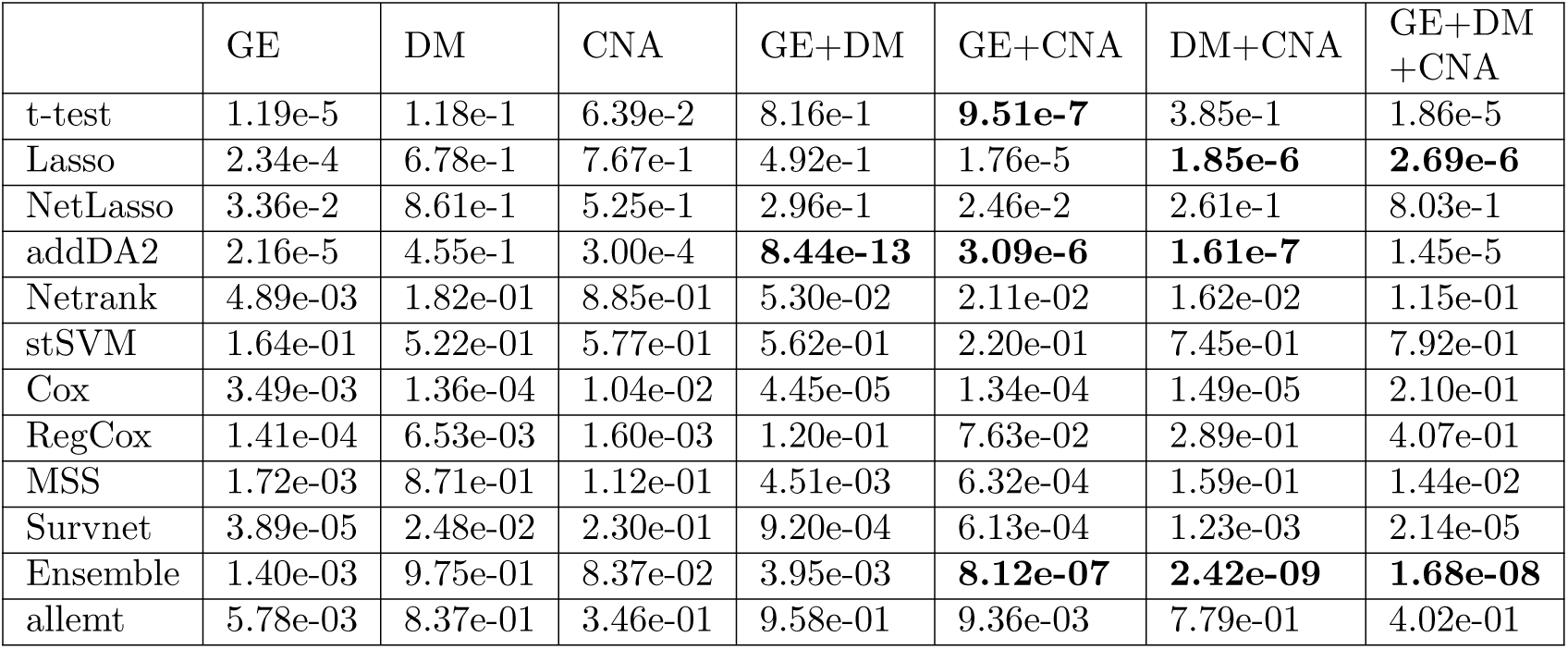
The p-values of log-rank tests based on the clustering of spectral clustering algorithm for different data level combinations using extended EMT network. We highlighted all p-values that are lower than 10e-5.

**Table S3.**
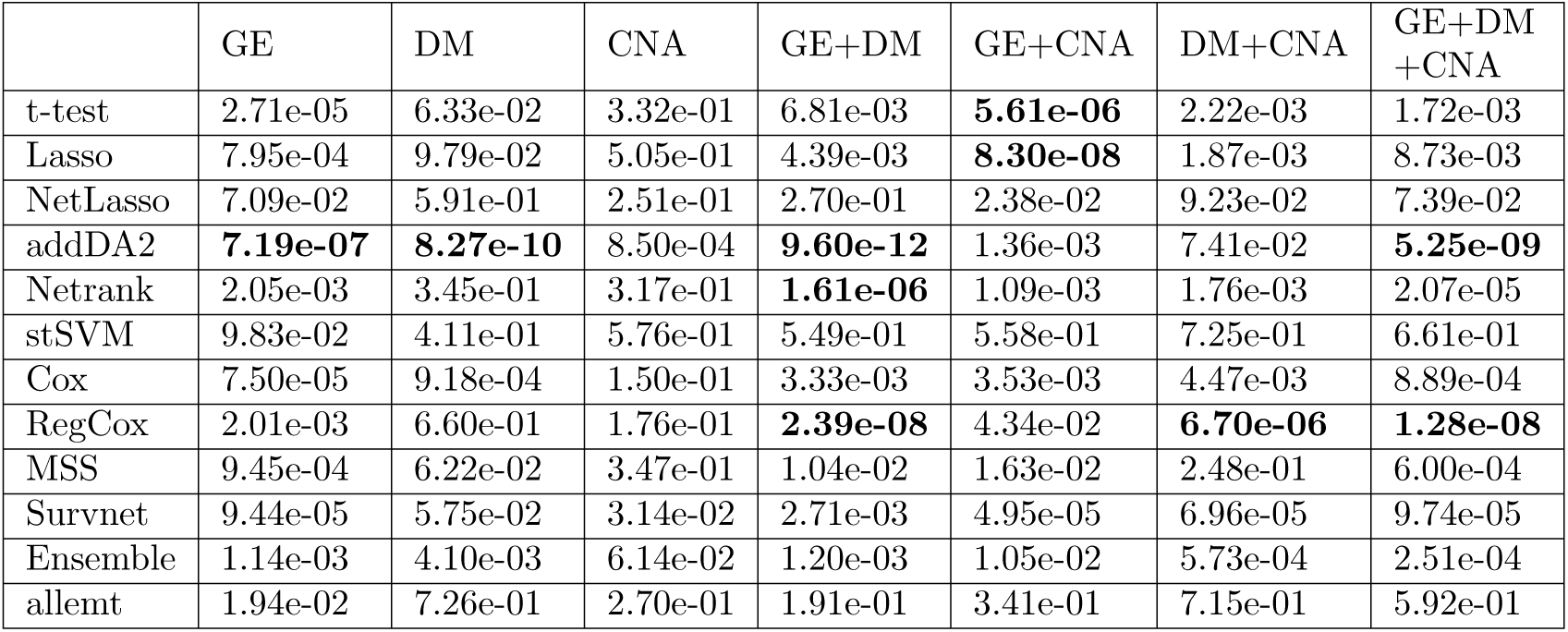
The p-values of log-rank tests based on SNF clustering using different data level combinations with extended EMT network. We highlighted all p-values that are lower than 10e-5.

**Table S4.**
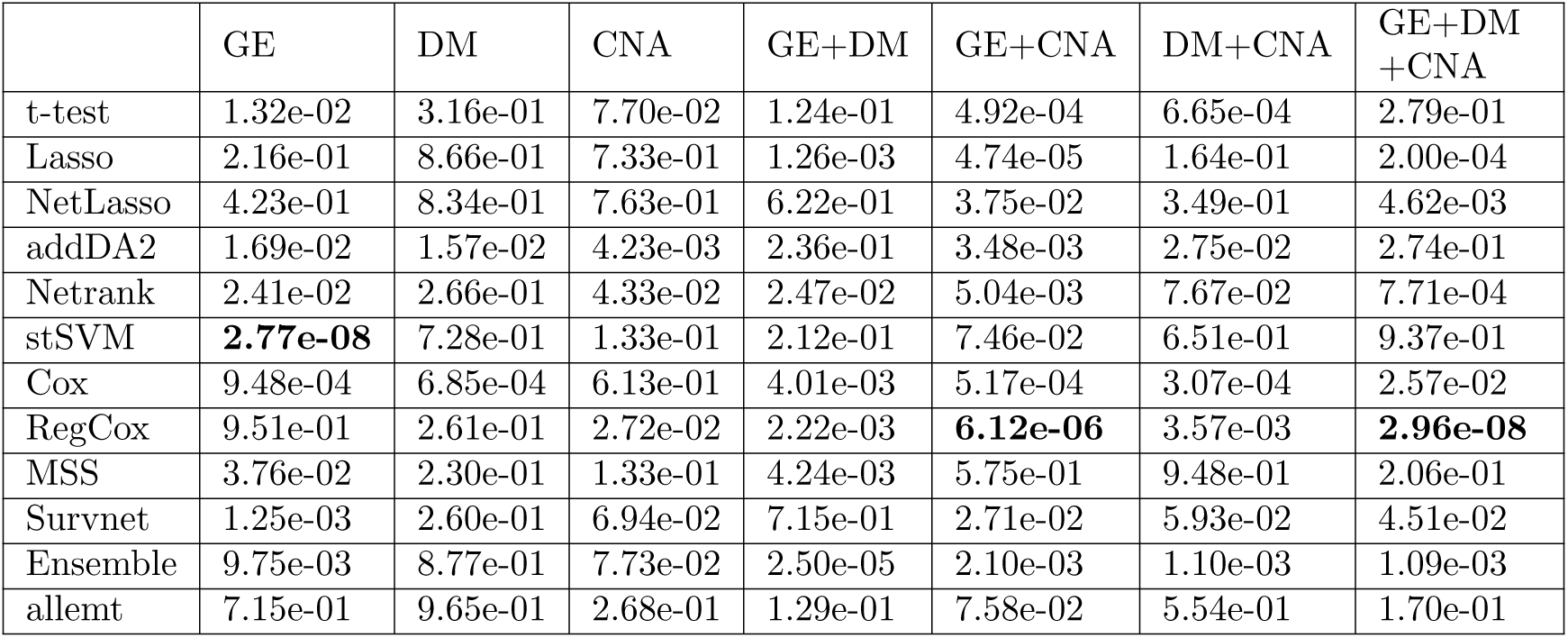
The p-values of log-rank tests based on iCluster clustering using different data level combinations with extended EMT network. We highlighted all p-values that are lower than 10e-5.

## References

1. Howlader N, Mariotto AB, Woloshin S, Schwartz LM. Providing clinicians and patients with actual prognosis: cancer in the context of competing causes of death. Journal of the National Cancer Institute Monographs. 2014;2014(49):255–264.

2. Subramanian J, Simon R. Gene expression–based prognostic signatures in lung cancer: ready for clinical use? Journal of the National Cancer Institute. 2010;102(7):464–474.

3. Lapointe J, Li C, Higgins JP, Van De Rijn M, Bair E, Montgomery K, et al. Gene expression profiling identifies clinically relevant subtypes of prostate cancer. Proceedings of the National Academy of Sciences of the United States of America. 2004;101(3):811–816.

4. Sørlie T, Perou CM, Tibshirani R, Aas T, Geisler S, Johnsen H, et al. Gene expression patterns of breast carcinomas distinguish tumor subclasses with clinical implications. Proceedings of the National Academy of Sciences. 2001;98(19):10869–10874.

5. Weinstein JN, Collisson EA, Mills GB, Shaw KRM, Ozenberger BA, Ellrott K, et al. The cancer genome atlas pan-cancer analysis project. Nature genetics. 2013;45(10):1113–1120.

6. Inza I, Larrãnaga P, Blanco R, Cerrolaza AJ. Filter versus wrapper gene selection approaches in DNA microarray domains. Artificial intelligence in medicine. 2004;31(2):91–103.

7. Dίaz-Uriarte R, De Andres SA. Gene selection and classification of microarray data using random forest. BMC bioinformatics. 2006;7(1):3.

8. Chuang HY, Lee E, Liu YT, Lee D, Ideker T. Network-based classification of breast cancer metastasis. Molecular systems biology. 2007;3(1).

9. Li J, Roebuck P, Grünewald S, Liang H. SurvNet: a web server for identifying network-based biomarkers that most correlate with patient survival data. Nucleic acids research. 2012; p. gks386.

10. Martinez-Ledesma E, Verhaak RG, Trevinõ V. Identification of a multi-cancer gene expression biomarker for cancer clinical outcomes using a network-based algorithm. Scientific reports. 2015;5.

11. Patel VN, Gokulrangan G, Chowdhury SA, Chen Y, Sloan AE, Koyutürk M, et al. Network Signatures of Survival in Glioblastoma Multiforme. PLoS Comput Biol. 2013;9(9):e1003237.

12. Allahyar A, de Ridder J. FERAL: network-based classifier with application to breast cancer outcome prediction. Bioinformatics. 2015;31(12):i311–i319.

13. Tang J, Alelyani S, Liu H. Feature selection for classification: A review. Data Classification: Algorithms and Applications. 2014; p. 37.

14. Yang S, Yuan L, Lai YC, Shen X, Wonka P, Ye J. Feature grouping and selection over an undirected graph. In: Graph Embedding for Pattern Analysis. Springer; 2013. p. 27–43.

15. Li C, Li H. Network-constrained regularization and variable selection for analysis of genomic data. Bioinformatics. 2008;24(9):1175–1182.

16. Zhang W, Ota T, Shridhar V, Chien J, Wu B, Kuang R. Network-based survival analysis reveals subnetwork signatures for predicting outcomes of ovarian cancer treatment. PLoS Comput Biol. 2013;9(3):e1002975.

17. Chen L, Xuan J, Riggins RB, Clarke R, Wang Y. Identifying cancer biomarkers by network-constrained support vector machines. BMC systems biology. 2011;5(1):161.

18. Morrison JL, Breitling R, Higham DJ, Gilbert DR. GeneRank: using search engine technology for the analysis of microarray experiments. BMC bioinformatics. 2005;6(1):233.

19. Winter C, Kristiansen G, Kersting S, Roy J, Aust D, Knösel T, et al. Google goes cancer: improving outcome prediction for cancer patients by network-based ranking of marker genes. PLoS Comput Biol. 2012;8(5):e1002511.

20. Cun Y, Fröhlich H. Network and data integration for biomarker signature discovery via network smoothed t-statistics. PLoS one. 2013;8(9):e73074.

21. Komurov K, Dursun S, Erdin S, Ram PT. NetWalker: a contextual network analysis tool for functional genomics. BMC genomics. 2012;13(1):282.

22. Cun Y, Fröhlich H. Prognostic gene signatures for patient stratification in breast cancer-accuracy, stability and interpretability of gene selection approaches using prior knowledge on protein-protein interactions. BMC bioinformatics. 2012;13(1):69.

23. Staiger C, Cadot S, Kooter R, Dittrich M, Müller T, Klau GW, et al. A critical evaluation of network and pathway-based classifiers for outcome prediction in breast cancer. PLoS one. 2012;7(4):e34796.

24. Staiger C, Cadot S, Györffy B, Wessels LF, Klau GW. Current composite-feature classification methods do not outperform simple single-genes classifiers in breast cancer prognosis. Frontiers in genetics. 2013;4.

25. Dao P, Colak R, Salari R, Moser F, Davicioni E, Schönhuth A, et al. Inferring cancer subnetwork markers using density-constrained biclustering. Bioinformatics. 2010;26(18):i625–i631.

26. Jin N, Wu H, Miao Z, Huang Y, Hu Y, Bi X, et al. Network-based survival-associated module biomarker and its crosstalk with cell death genes in ovarian cancer. Scientific reports. 2015;5.

27. Gwinner F, Boulday G, Vandiedonck C, Arnould M, Cardoso C, Nikolayeva I, et al. Network-based analysis of omics data: The LEAN method. Bioinformatics. 2016; p. btw676.

28. Seoane JA, Day IN, Gaunt TR, Campbell C. A pathway-based data integration framework for prediction of disease progression. Bioinformatics. 2014;30(6):838–845.

29. Mounika Inavolu S, Renbarger J, Radovich M, Vasudevaraja V, Kinnebrew G, Zhang S, et al. IODNE: An integrated optimization method for identifying the deregulated subnetwork for precision medicine in cancer. CPT: Pharmacometrics & Systems Pharmacology. 2017;.

30. Ideker T, Ozier O, Schwikowski B, Siegel AF. Discovering regulatory and signalling circuits in molecular interaction networks. Bioinformatics. 2002;18(suppl 1):S233–S240.

31. Zhang Y CRRH Xuan J. Module-Based Breast Cancer Classification. International Journal of Data Mining and Bioinformatics. 2013;7:284–302.

32. Wu G, Stein L. A network module-based method for identifying cancer prognostic signatures. Genome biology. 2012;13(12):R112.

33. Ein-Dor L, Zuk O, Domany E. Thousands of samples are needed to generate a robust gene list for predicting outcome in cancer. Proceedings of the National Academy of Sciences. 2006;103(15):5923–5928.

34. Michiels S, Koscielny S, Hill C. Prediction of cancer outcome with microarrays: a multiple random validation strategy. The Lancet. 2005;365(9458):488–492.

35. Ein-Dor L, Kela I, Getz G, Givol D, Domany E. Outcome signature genes in breast cancer: is there a unique set? Bioinformatics. 2004;21(2):171–178.

36. Venet D, Dumont JE, Detours V. Most random gene expression signatures are significantly associated with breast cancer outcome. PLoS Comput Biol. 2011;7(10):e1002240.

37. Shao B. Phenotype Relevant Network-based Biomarker Discovery Integrating Multiple Omics Data. Freie Universität Berlin; 2018.

38. Xia J, Benner MJ W Hancock RE. NetworkAnalyst - integrative approaches for protein–protein interaction network analysis and visual exploration. Nucleic Acids Research. 2014;.

39. Shao B, Cannistraci CV, Conrad TO. Epithelial Mesenchymal Transition Network-Based Feature Engineering in Lung Adenocarcinoma Prognosis Prediction Using Multiple Omic Data. Genomics and Computational Biology. 2017;3(3):57.

40. Haury AC, Gestraud P, Vert JP. The influence of feature selection methods on accuracy, stability and interpretability of molecular signatures. PLoS one. 2011;6(12):e28210.

41. Tibshirani R. Regression shrinkage and selection via the lasso. Journal of the Royal Statistical Society Series B (Methodological). 1996; p. 267–288.

42. Cox DR. Regression Models and Life-Tables. Journal of the Royal Statistical Society Series B (Methodological). 1972;34(2):187–220.

43. Network CGAR, et al. Integrated genomic analyses of ovarian carcinoma. Nature. 2011;474(7353):609–615.

44. Simon N, Friedman J, Hastie T, Tibshirani R. Regularization paths for Cox’s proportional hazards model via coordinate descent. Journal of statistical software. 2011;39(5):1.

45. Li J, Lenferink AE, Deng Y, Collins C, Cui Q, Purisima EO, et al. Identification of high-quality cancer prognostic markers and metastasis network modules. Nature communications. 2010;1:34.

46. Thiery JP. Epithelial–mesenchymal transitions in tumour progression. Nature Reviews Cancer. 2002;2(6):442–454.

47. Thiery JP, Acloque H, Huang RY, Nieto MA. Epithelial-mesenchymal transitions in development and disease. cell. 2009;139(5):871–890.

48. Hanahan D, Weinberg RA. Hallmarks of cancer: the next generation. cell. 2011;144(5):646–674.

49. Schliekelman MJ, Taguchi A, Zhu J, Dai X, Rodriguez J, Celiktas M, et al. Molecular portraits of epithelial, mesenchymal, and hybrid States in lung adenocarcinoma and their relevance to survival. Cancer research. 2015;75(9):1789–1800.

50. Brabletz T, Kalluri R, Nieto MA, Weinberg RA. EMT in cancer. Nature Reviews Cancer. 2018;.

51. Hahsler M, Grün B, Hornik K. A computational environment for mining association rules and frequent item sets. 2005;.

52. Agrawal R, Imieliński T, Swami A. Mining association rules between sets of items in large databases. In: Acm sigmod record. vol. 22. ACM; 1993. p. 207–216.

53. Agrawal R, Srikant R, et al. Fast algorithms for mining association rules. In: Proc. 20th int. conf. very large data bases, VLDB. vol. 1215; 1994. p. 487–499.

54. Wang B, Mezlini AM, Demir F, Fiume M, Tu Z, Brudno M, et al. Similarity network fusion for aggregating data types on a genomic scale. Nature methods. 2014;11(3):333–337.

55. Shen R, Olshen AB, Ladanyi M. Integrative clustering of multiple genomic data types using a joint latent variable model with application to breast and lung cancer subtype analysis. Bioinformatics. 2009;25(22):2906–2912.

56. Bjaanæs MM, Fleischer T, Halvorsen AR, Daunay A, Busato F, Solberg S, et al. Genome-wide DNA methylation analyses in lung adenocarcinomas: association with EGFR, KRAS and TP53 mutation status, gene expression and prognosis. Molecular oncology. 2016;10(2):330–343.

57. Shao B, Conrad T. Epithelial-Mesenchymal Transition Regulatory Network-Based Feature Selection in Lung Cancer Prognosis Prediction. In: International Conference on Bioinformatics and Biomedical Engineering. Springer; 2016. p. 135–146.

58. Kim D, Li R, Lucas A, Verma SS, Dudek SM, Ritchie MD. Using knowledge-driven genomic interactions for multi-omics data analysis: metadimensional models for predicting clinical outcomes in ovarian carcinoma. Journal of the American Medical Informatics Association. 2016;24(3):577–587.

59. Huang HL, Wu YC, Su LJ, Huang YJ, Charoenkwan P, Chen WL, et al. Discovery of prognostic biomarkers for predicting lung cancer metastasis using microarray and survival data. BMC Bioinformatics. 2015;16(1).

60. Futreal PA, Coin L, Marshall M, Down T, Hubbard T, Wooster R, et al. A census of human cancer genes. Nature Reviews Cancer. 2004;4(3):177–183.

61. Chikaishi Y, Uramoto H, Tanaka F. The EMT status in the primary tumor does not predict postoperative recurrence or disease-free survival in lung adenocarcinoma. Anticancer research. 2011;31(12):4451–4456.

62. Zhao J, Dong D, Sun L, Zhang G, Sun L. Prognostic significance of the epithelial-to-mesenchymal transition markers e-cadherin, vimentin and twist in bladder cancer. International braz j urol. 2014;40(2):179–189.

